# Contemporary hybridization among *Arabis* floodplain species creates opportunities for adaptation

**DOI:** 10.1101/2025.07.03.662201

**Authors:** Neda Rahnamae, Lukas Metzger, Lea Hördemann, Kevin Korfmann, Abdul Saboor Khan, Craig I. Dent, Samija Amar, Raúl Y. Wijfjes, Tahir Ali, Gregor Schmitz, Benjamin Stich, Aurelien Tellier, Juliette de Meaux

**Affiliations:** Institute for Plant Sciences, Biocenter, Cologne, Germany; Institute of Plant Ecology and Evolution, Heinrich-Heine University Düsseldorf, Düsseldorf, Germany; Professorship for Population Genetics, Department of Life Science Systems, Technical University of Munich, Freising, Germany; Research Department for Limnology, Mondsee, University of Innsbruck, Innsbruck, Austria; Institute of Ecology and Evolution, University of Oregon, Eugene, OR, USA; Department of Chromosome Biology, Max Planck Institute for Plant Breeding Research, Cologne, Germany; CEPLAS: Cluster of Excellence on Plant Sciences, Heinrich-Heine University, Düsseldorf, Germany; Faculty of Biology, LMU Munich, Planegg-Martinsried, Germany; Julius Kühn Institute (JKI), Federal Research Centre for Cultivated Plants, Institute for Breeding Research on Agricultural Crops, Sanitz, Germany

**Keywords:** hybridization, QTL mapping, meiotic drive, heterozygote advantage, competitive grasslands, *Arabis* genome

## Abstract

Hybridization between closely related species is increasingly recognized as a major source of biodiversity. Yet, whether it can create advantageous trait combinations while purging harmful alleles remains unknown. We studied *Arabis nemorensis* and *A. sagittata*, two endangered species that currently hybridize in a single hotspot. We measured 22 phenotypic traits and mapped their genetic basis in an F2 population, after generating high quality genome assemblies for both species. In total, 58 QTLs were identified for 20 traits, with additive and dominance effects best fitting Gaussian and logistic distributions, respectively. Six large-effect QTLs were linked to significant hybrid fitness loss. Two genomic regions showed strong transmission bias favoring *A. sagittata* alleles, potentially accelerating their introgression. However, 48% of QTLs were unlinked to reduced fitness or segregation distortion and may generate genotypes exceeding parental performance. Notably, a major QTL affecting flowering time explained 23% of phenotypic variation and implicated *TFL1* as a candidate gene for life history adaptation. While most QTLs lacked overlap with past selective sweeps, indicating limited recent positive selection, 5 of 7 QTLs for rosette size overlapped with sweep signatures in the parental lineages. Overall, our findings offer unique insights into incipient stages of hybridization.

## Introduction

Biodiversity is crucial for the structure and functioning of ecosystems (Staudinger *et al*., 2012). The direct impact of human activity and the escalating threat of climate change have initiated the sixth Mass Extinction (Cochrane *et al*., 2016; Ceballos *et al*., 2020; Cowie *et al*., 2022; IPCC, 2023), sparking global concern (Hooper *et al*., 2002; Eichenberg *et al*., 2021; Theissinger *et al*., 2023) and prompting investigation into how species will cope (Moore & Hendry, 2009; Bontrager & Angert, 2019; Schlaepfer & Lawler, 2023). For species endangered by rapid shifts in the environment, the acquisition of pre-adapted alleles via hybridization could provide plants with a strategy to immediately combat climate change (Hansen *et al*., 2012; Anderson *et al*., 2012; Chunco, 2014; Brauer *et al*., 2023).

Hybridization, the interbreeding of individuals from genetically distinct populations or species, occurs frequently among close relatives (Blanckaert *et al*., 2023; Peñalba *et al*., 2024; Rosser *et al*., 2024). It can have both advantageous and detrimental consequences on the species receiving gene flow (Peñalba *et al*., 2024). First, interspecific hybridization can enable locally adapted alleles across species barriers to be transferred and thus enhance the adaptive potential of species (Seehausen, 2004; Pfennig *et al*., 2016; Abbott, 2017). Indeed, hybridization between populations with different ecological specializations can give rise to new, viable, and fertile hybrids equipped with novel trait combinations. Such combinations may improve the fitness of the population or even enable previously untapped habitats to be colonized (Buerkle *et al*., 2000; Rieseberg *et al*., 2003; Mallet, 2007; Abbott *et al*., 2013; Blanckaert *et al*., 2023). Because this process can contribute to the rescue of endangered species, it may help maintain diversity, as seen in genus *Pachyclasdon*, which appears to have survived the Last Glacial Maximum thanks to genetic information transferred through hybridization in alpine refugia on New Zealand’s South Island (Becker *et al*., 2013). Early-flowering alleles of *Helianthus argophyllus* were enriched among locally adapted introgressed genomic fragments in *Helianthus* spp and *Betula platyphtylla* (Todesco *et al*., 2020; Nocchi *et al*., 2023). Hybrid populations between generalists and narrow range endemic rainbowfishes (*Melanotaenia* spp.) were well adapted to environmental changes thanks to the effects of introgression (Brauer *et al*., 2023). Hybridization has even been proposed to promote the formation of new species, a phenomenon referred to as “hybrid speciation”; however, whether a new species actually results is controversial (Anderson & Stebbins, 1954; Ellstrand & Schierenbeck, 2000; Schumer *et al*., 2015).

The detrimental effects of hybridization are often more readily detected than its advantageous effects. Indeed, allelic incompatibilities can cause a massive fitness breakdown, when gene pools reunite after long periods of evolution in isolation (e.g., *Drosophila* [Masly & Presgraves, 2007; Cooper *et al*., 2018]; *Mimulus* [Zuellig & Sweigart, 2018]; *Mus* [White *et al*., 2011; White *et al*., 2012; Wang *et al*., 2015]; *Xiphophorus* [Schumer *et al*., 2014; Schumer & Brandvain, 2016; Powell *et al*., 2020; Moran *et al*., 2024]; Bomblies & Weigel, 2007; Presgraves, 2010; Coughlan & Matute, 2020; Li *et al*., 2022). The overall fitness of the hybridizing population will be reduced if too many resources are used to produce poorly performing hybrids (Rhymer & Simberloff, 1996; Todesco *et al*., 2016; Goulet *et al*., 2017). This phenomenon, sometimes described as “demographic swamping,” elevates the risk of extinction but may also select for allelic variations that reinforce species isolation (Hopkins, 2013; Goulet *et al*., 2017; Ma *et al*., 2019; Brauer *et al*., 2023).

Despite the interest in the positive consequences of hybridization and the abundant evidence for allelic incompatibilities (Bomblies & Weigel, 2007), we know little about how the positive and negative effects of hybridization interact, much less about how a genotype carrying adaptive alleles might emerge – especially against a background in which detrimental effects have been recombined out. Such “super genotypes,” although rare in the offspring of the first generation of hybrids (many of which may perform poorly), may represent new and exceptional combinations of adaptive alleles that in selfing species can determine the evolutionary success of hybridization. In addition, much focus has been placed on showing that introgressed alleles bring adaptive advantages to the recipient species (Edelman and Mallet 2021). Yet, the role that natural selection has played in shaping the introgressing alleles of the donor species has seldom been examined. Addressing these two challenges is best achieved when hybridization can be studied as it proceeds, as this allows direct observation of the recombination and selection dynamics shaping introgressed variation.

Here, we focused on a hybridization hotspot located along the banks of the Rhine River near Mainz, Germany, where two endangered Arabis species hybridize in a floodplain meadow (Dittberner *et al*., 2019; Dittberner *et al*., 2022). *Arabis nemorensis*, a species within the Brassicaceae family (*Arabis hirsuta* tribe), inhabits floodplain meadows and is currently in a critical state in Central Europe, requiring special attention from conservation authorities (Schnittler & Günther, 1999; Burmeier *et al*., 2011). *A. nemorensis* is self-pollinating and exhibits low levels of nucleotide diversity. Its endangered status is further intensified by its unique ecological requirements, and the loss of its natural habitat (Hölzel, 2005; Burmeier *et al*., 2011; Mathar *et al*., 2015; Dittberner *et al*., 2019). *A. sagittata*, another member of the same phylogenetic tribe (Karl & Koch, 2014), is morphologically very similar but commonly found in calcareous grasslands and thus possibly more tolerant to drought (Dittberner *et al*., 2022). *A. sagittata* was recently observed in floodplains, where it naturally hybridizes with *A. nemorensis* (Dittberner *et al*., 2019; Dittberner *et al*., 2022).

Introgression analysis has revealed that gene flow between *A. nemorensis* and *A. sagittata* has happened in the past, but contemporary hybridization is restricted to the sympatric population, where intraspecific genetic variation is extremely low (Dittberner *et al*., 2019; Dittberner *et al*., 2022). Additionally, population genetics analyses estimated the divergence between *A. nemorensis* and *A. sagittata* occurred around 900,000 generations ago, with practically complete isolation after the last glaciation (Dittberner *et al*., 2022).

Using the F2 progeny of a cross between two representative genotypes of *A. nemorensis* and *A. sagittata* from the hybridizing hotspot, we addressed the following questions: (1) What are the phenotypic differences of relevant ecological traits? (2) What is the genetic architecture of interspecific differences in this hybridizing hotspot? (3) Can a new combination of ecologically relevant traits arise and establish after hybridization? (4) Do QTL regions enriched in genomic regions carry footprints of past selection? Our study confirmed the extent of the genetic differences underpinning phenotypic divergence between two species. In addition, we found that some F2 hybrids show extreme trait values, despite their generally lower seed production compared to parental lines. Interestingly, incompatibility QTLs have a simple genetic basis, and some ecologically relevant QTLs are independent from the incompatibility QTLs. Collectively, these results indicate that a few offspring of hybridized species may potentially harbor a genotype with a combination of properties that none of the parents have.

## Materials and Methods

### Common Garden Experiment and Phenotyping

We crossed sympatric *A. nemorensis* genotype 10 with the *A. sagittata,* genotype 69, collected from the banks of the Rhine River near Mainz in Riedstadt, Hessen, Germany in 2015 and fully sequenced (Dittberner *et al*., 2019; Dittberner *et al*., 2022). Because nucleotide diversity within the population is very low (*A. nemorensis* π = 4.37e-5 and *A. sagittata* π = 1.32e-5, synonymous sites, Dittberner *et al*., 2022), we assume here that differences between these genotypes will predominantly reflect differences between species. Plants were reciprocally crossed to generate F1s, and we noted that their fitness was comparable to that of the parental species (Fig. S1). Seedlings were grown in the greenhouse of the Experimental Garden of the University of Cologne, and the seeds of the first generation of selfing (F2) were harvested. We sowed F2 seeds in trays on 05.10.2019, and after two weeks of vernalization and germination we transplanted seedlings into 7x7cm pots filled with *Topferde* soil (Einheitserde, Sinntal-Altengronau, Germany), placing one seedling per pot. In total, 1,204 individual plants (both hybrids and parental replicates) were distributed across 43 trays, which were put in cold frames to accelerate growth and prevent frost damage from 08.11 to 13.12.2018. On 13.12.2018, we transferred trays to bird-protected cages under seminatural conditions. On 19.11.2018, photographs of each tray were taken using a Canon EOS and rosette sizes were measured using ImageJ. Also, on 19.12.2018, we started harvesting leaves from plants that were big enough (∼50 mg of leaves per plant). Harvesting continued in January, February, and March 2019. Harvested rosette leaves were stored at -80°C for DNA extraction.

On 01.03.2019, eight trays were moved to cold frames for a 10-day period to recover from potential stress. A total of 199 F2 individual plants, along with seven *A. nemorensis* and seven *A. sagittata*, were submerged in transparent boxes, each containing 17 L of water for seven weeks. Plants were randomized and distributed across eight boxes, which were kept in cold frames throughout the submergence experiment. After seven weeks of complete submergence, plants were removed from water and left to recover. Then, their survival status was documented. Following a two-week recovery period, we again recorded the survival status of plants, categorizing plants as either dead, survived with new leaf growth, or survived and bolted. During recovery, pots remained in cold frames and were watered as necessary.

Between November 2018 and April 2019, we recorded more than 20 phenotypic traits for each plant. These traits were grouped into five categories: (1) fitness: seed mass from 10 randomly selected siliques or fertility score and survival status after submergence; (2) growth: rosette diameter at four time points (19.11.2018, 08.01.2019, 07.02.2019, 13.03.2019), inflorescence height, and ena of side and ground shoots; (3) timing: bolting and flowering times; (4) rosette leaf Ttraits: stem leaf number, leaf length, petiole length, lamina length, lamina length/width ratio, leaf margin type (serrated or smooth), leaf width, and petal length; and (5) stem traits: stem leaf density, stem leaf length and width, and stem height.

### Phenotypic Analyses

To assess phenotypic differences between the two species, we used a generalized linear model (trait ∼ Tray, with quasipoisson distribution of error) to analyze each trait, controlling for experimental effects, in R. A t-test from the stats package in R (version 3.6.2) was then applied to the residuals of the models to calculate the significance of differences (*p*-value < 0.05) between the two groups (*A. nemorensis* and *A. sagittata*, each with 35 replicates) for each trait. We used ggplot2 (version 3.5.1; Wickham, 2011) to visualize the distribution of phenotypes for both F2 and parental replicates in a single plot per trait, and to inform our understanding of transgressive segregation within the F2 population (Suppl. File 1).

For genetically based phenotypic correlation between individuals of the F2 population, we employed a mixed-effects model approach (trait ∼ Cross/TrayBlock) that accounted for random variation attributable to experimental blocks and trays, and cross direction due to reciprocal cross. Residuals were extracted for each trait, and pairwise Spearman correlations were calculated using the *corr.test* function from the psych package in R (version 2.4.6.26); pairwise deletions were applied to handle missing data. Correlation significance was assessed at α = 0.05. The resulting correlation and *p*-value matrices were used to construct a network in which only significant correlations were retained. A network graph was generated using the igraph (version 2.1.1) and ggraph (version 2.1.1) packages. Each node represents a phenotypic trait, and edges represent statistically significant pairwise correlations. Edge color indicates the sign of the correlation (green for positive, brown for negative), and edge thickness is scaled to the absolute strength of the correlation. Curved edges and partial transparency were used to enhance visual clarity, and node labels were plotted within stylized trait circles. The *ggcorrplot* function from the ggcorrplot package (version 0.1.4.1) was used to visualize the heatmap.

### DNA Extraction and RAD-seq Library Construction

DNA was extracted from leaves stored at -80°C using the Nucleospin® 8 Plant II protocol. Using the restriction-site-associated DNA sequencing (RAD-seq) protocol described in Dittberner *et al*. (2019), we analyzed the genomes of 801 F2 individuals. DNA was quantified with Qubit® 3.0. Fluorometer. For each sample, a total of 250 ng of genomic DNA was digested with the KpnI-HF restriction enzyme (New England Biolabs). Each digested sample was then ligated with one individually barcoded modified Illumina P1 adapter containing 5 bp random nucleotides to PCR duplicates to be removed in the steps that follow. Twenty different barcodes were used, and 52 pools of 20 barcoded individuals each were constructed, with all pools standardized to equal volume and concentration. Libraries were sequenced in four separate runs at the Cologne Center for Genomics (CCG) on the Illumina NovaSeq platform. For 46 pools, sequencing length was 2x100 bp, generating approximately 3 million reads per individual; for the remaining 6 pools, sequencing length was 2x150 bp, generating approximately 6 million reads per individual. These RADseq libraries covered, on average, approximately 2% of the 248 Mb genome.

### Genome Assembly and Annotation

In order to improve previous genome assemblies, the DNA of both parental lines was sequenced with PacBio HiFi technology and Hi-C data. We used Jellyfish (version 2.3.0; Marçais & Kingsford, 2011) to count k-mers of size 21 in the 2.18 and 1.98 million reads obtained for *A. nemorensis* and *A. sagittata*, respectively. The k-mer histogram generated by Jellyfish was then processed with GenomeScope (version 2.0; Ranallo-Benavidez *et al*., 2020) to estimate genome size, heterozygosity, and repetitiveness. HiFiAdapterFilt (version 2.0.0; Sim *et al*., 2022) was used to remove residue PacBio adapter sequences from the HiFi reads. Then, hifiasm was used again to assemble the filtered HiFi reads (version 0.16.1; Cheng *et al*., 2021; Cheng *et al*., 2022) with integration of Hi-C data.

Hi-C reads were aligned to the draft contigs using BWA (version 0.7.17; Li & Durbin, 2009), and the resulting alignments were processed with the Juicer pipeline (version 1.6; Durand *et al*., 2016b) to generate chromatin contact maps. These maps were manually inspected and curated using Juicebox (version 2.20.00; Durand *et al*., 2016a) to correct misjoins and to anchor and orient contigs into chromosome-scale scaffolds. This integration significantly enhanced the contiguity and accuracy of the assembly.

To scaffold the primary assembly produced by hifiasm, we employed RagTag (version 2.1.0; Alonge *et al*., 2022), using the *A. alpina* assembly as a reference (Suppl. File 2) to further improve scaffold ordering and correct structural inconsistencies. In addition, a linkage map (see below) allowed us to correct a few minor assembly errors. The final assemblies have been deposited in the European Nucleotide Archive (ENA) and is awaiting an accession number (Project id: PRJEB89863). The ab-initio annotation was generated using the *A. nemorensis* reference genome, which was uploaded to the PlabiPD Helixer platform (https://www.plabipd.de/helixer_main.html). The annotation was performed in “Lineage-Specific” mode with selection of the option “Land Plant”.

### Analysis of RAD-seq Data, SNP Calling, and Construction of Linkage Map

RAD-seq read quality was assessed with FastQC (version 0.11.9; Andrews, 2010). PCR duplicates were removed using the clone_filter module in Stacks (version 2.59; Catchen *et al*., 2013), based on a 5 bp random nucleotide sequence at the end of each adapter. Adapters were trimmed and reads shorter than 60 bp were removed using Cutadapt (version 1.18; Martin, 2011). We demultiplexed samples and filtered out reads with ambiguous barcodes (allowing one mismatch), cut sites, uncalled bases, and low-quality reads (default threshold) using the process_radtags module in Stacks (version 2.59). Reference-based mapping and read-filtering were conducted with BWA (version 0.7.17; Li & Durbin, 2009) using default settings against the *A. nemorensis* reference genome, along with SAMtools (version 1.10; Li *et al*., 2009) and bash scripts from Rivera-Colón & Catchen (2022) (Suppl. File 3).

We performed variant-calling for 801 individuals using BCFtools mpileup and call (version 1.18; Li *et al*., 2009) under specific criteria: a base quality score greater than 30, base quality recalculation (-E option), and SNP calling with a significance threshold of *p*-value < 0.05. Genotyped loci were filtered using VCFtools (version 0.1.17; Danecek *et al*., 2011) to exclude loci with more than 50% missing data across individuals. Only biallelic sites were kept, and indels were removed. Individuals with over 60% missing data were excluded, leaving a working set of 781 individuals for subsequent analysis. Additional filters were applied to site and genotype depth (--min-meanDP 4 --max-meanDP 40 --minDP 5 --maxDP 40), and sites with more than 70% missing data were then removed. Loci spaced less than 100 bp apart were clustered into RAD regions as described by Dittberner *et al*. (2019). Regions with abnormally high or low coverage were excluded based on specific thresholds: mean coverage greater than twice or less than one-third the overall mean, maximum coverage exceeding twice the mean maximum coverage across all regions, or regions shorter than 150 bp. Sites absent from parental lines and ambiguous bases were also removed. Finally, we applied a minor allele frequency filter (--maf 0.25) to the dataset. SNPs (single nucleotide polymorphisms) were extracted using VCFtools (version 0.1.17; Danecek *et al*., 2011) and custom Python scripts, resulting in a VCF file containing 5,360 SNP markers across 781 individuals (Suppl. File 3).

We applied additional filters to improve the quality of map construction. We removed 39 individuals with missing data exceeding 3,500 loci. Duplicated markers were identified and removed, resulting in 47 markers being discarded. Subsequently, markers with more than 23% missing data were filtered out. To analyze the remaining 2,164 markers, we examined their distribution along the genome, compared segregation distortion patterns with allelic proportion and missingness, and filtered out markers identified as outliers based on allelic proportion. We further removed markers with fractions of heterozygotes lower than 25% or higher than 75%. We also used RepeatMasker (version 4.1.6; Smit *et al*., 2015) on genome annotations to detect and remove markers within repetitive regions. We used repeatmodeler to identifiy repeat families, an approach that was used in Repeatmasker to mask the repetitive regions in the genome. This final analysis produced a genetic map of 2,082 markers distributed across eight chromosomes for 742 individuals (Suppl. File 4).

To improve the accuracy of genotype calling, particularly for heterozygous loci located in low-coverage regions, a Python script was developed to correct incorrectly called genotypes and impute missing data using a sliding window approach. The 2 Mb window size with a 0.5 Mb step size demonstrated the most effective correction and imputation. No further markers or individuals were excluded after this correction and imputation step. The finalized corrected dataset was then used to construct the genetic map using the ASMap package in the R environment (version 1.0-7; Taylor & Butler, 2017) packages in R (version 4.2.3), applying the so-called Kosambi mapping function in the R/qtl package (version 1.66; Broman *et al*., 2003) as detailed in Supplemental File 4.

Chromosomal orientation was assessed using the constructed genetic map, identifying inversions on chromosomes 3, 4, 6, and 7 due to assembly issues. These regions were adjusted by inverting physical distances on the genetic map, and the genome assembly was subsequently corrected. See above (Genome Assembly and Annotation) for details on the genome assembly process.

### Segregation Distortion and Selection Coefficient

We assessed segregation distortion within each region by applying a test of segregation distortion (profileMark) for each marker with Bonferroni correction using the ASMap package in R. To assess deviations from the Hardy-Weinberg equilibrium (HWE) and estimate selection coefficients, allele frequencies for each genotype (NN, NS, SS) were calculated based on the total number of individuals (N = 742). The observed genotype counts were used to compute allele frequencies (*Freq_N_* and *Freq_S_*) by summing homozygous and heterozygous contributions. Expected genotype frequencies under the HWE were then computed as *Freq_N_^2^* (NN), 2 × *Freq_N_* × *Freq_S_* (NS), and *Freq_S_^2^* (SS). These frequencies were scaled to expected counts by multiplying by the total number of individuals (Suppl. File 4).

To determine significant deviations from the HWE, a chi-squared goodness-of-fit test was performed for each locus. The observed and expected genotype counts were compared using the chi-squared statistic, with 2 degrees of freedom. Significant deviations were identified based on a Bonferroni-corrected *p*-value threshold, accounting for multiple testing across all loci. The Bonferroni correction was applied by adjusting the significance threshold to 0.05 divided by the total number of markers (2,082); this adjustment ensured that false positives were stringently controlled for. Additionally, selection coefficients (t and s) were estimated by calculating the relative differences in ratios of observed to expected frequencies among genotypes. Specifically, the coefficients were defined 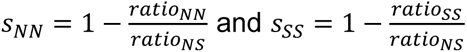 reflecting the relative fitness difference between homozygotes and heterozygotes (Carey & Ganders, 1980) (Suppl. File 4).

Significance of genotype deviations from the HWE was assessed using a chi-squared test for each marker. *P*-values were adjusted for multiple testing using the false discovery rate (FDR) correction method, controlling the FDR at 5%. Markers with FDR-adjusted *p*-values below 0.05 were considered significant and were visually highlighted in the selection coefficient plot. Red and blue circles denote significant selection against NN (t) and SS (s), respectively. When both t and s were significantly greater than zero, markers were interpreted as showing potential selection favoring the heterozygote (NS).

### QTL Mapping

QTL analyses were conducted using the R/qtl and QTLtools packages (version 1.66; Broman *et al*., 2003; version 1.3.1; Delaneau *et al*., 2017) in R. QTL mapping was performed on the residuals extracted from models that accounted for positional effects. Additionally, the effect of variation in the chloroplast and mitochondrial genomes was assessed by including cross direction in the model alongside environmental effects. QTL mapping was conducted for 22 traits listed in Table 1. However, survival after flooding, being a binomial trait, lacked a sufficient number of F2 individuals for QTL mapping (Suppl. File 5).

**Table 1.** Differences in phenotypic traits between parents. This table summarizes the phenotypic traits measured in the common garden experiment for F2 hybrids and their parental replicates (*A. nemorensis* and *A. sagittata*). Means represent the average value for each trait within parental replicates (35 individuals per species). “Distance” shows the absolute difference between the species’ means. The *p*-values indicate the significance of the difference between species based on t-statistics.

| Trait | Mean *A. nem* | Mean *A. sag* | Distance | $p$-value |
| --- | --- | --- | --- | --- |
| Days to Bolting (B.T) | 157.2857 | 169.2857 | 12 | 1.302E-19 |
| Days to Flowering (F.T) | 183.5714 | 193.4286 | 9.8571 | 5.556E-14 |
| Fertility Score (W.S) | 0.0298 | 0.0295 | 0.0004 | >0.63 |
| Inflorescence Height (P.H) | 35.2500 | 34.5714 | 0.6786 | 4.537E-02 |
| Lamina Length (Lam) | 1.2729 | 1.4646 | 0.1916 | 1.179E-03 |
| Lamina L/W (LLW) | 1.4760 | 1.7938 | 0.3178 | 3.702E-12 |
| Leaf Length (L.L) | 1.5562 | 1.7161 | 0.1599 | >0.75 |
| Leaf Width (L.W) | 0.8589 | 0.8213 | 0.0376 | 1.667E-02 |
| Number of Stem Leaves (N.L) | 28.1429 | 18.1429 | 10 | 4.567E-20 |
| Petal Length (Pet) | 4.0569 | 5.3579 | 1.3010 | 1.887E-21 |
| Petiole Length (Pti) | 0.2833 | 0.2516 | 0.0317 | 1.084E-06 |
| Rosette Diameter 1 (RD1) | 1.8331 | 2.2419 | 0.4087 | >0.23 |
| Rosette Diameter 2 (RD2) | 3.2946 | 3.6893 | 0.3947 | 4.922E-03 |
| Rosette Diameter 3 (RD3) | 3.3611 | 3.4424 | 0.0813 | 1.502E-03 |
| Rosette Diameter 4 (RD4) | 3.8805 | 3.9110 | 0.0305 | 7.447E-03 |
| Side Shoots (Ssh) | 2.7857 | 2.1429 | 0.6429 | 2.802E-02 |
| Stem Height (S.H) | 40.4286 | 34.4286 | 6 | 2.668E-09 |
| Stem Leaf Density (SLD) | 0.6979 | 0.5271 | 0.1707 | 5.508E-16 |
| Stem Leaf Length (SLL) | 2.3894 | 2.7921 | 0.4028 | 1.684E-04 |
| Stem Leaf Width (SLW) | 1.4021 | 1.2337 | 0.1684 | 2.966E-03 |
| Petiole L/Lamina L (P.L) | 0.2464 | 0.1886 | 0.0579 | 5.248E-07 |
| Ground Shoots (Gsh) | 0.2143 | 1.2857 | 1.0714 | 1.414E-08 |

For each trait, we used the *scantwo* function to perform a two-dimensional genome scan with a two-QTL model, applying the Haley-Knott regression algorithm (Haley & Knott, 1992). Penalties were calculated from 1,000 permutations of the *scantwo* function to support the stepwise fitting of multiple QTL models. The *stepwiseqtl* function was then employed, with a maximum of five QTL, using Haley-Knott regression to search for optimized models. We activated the *refine.locations* option in *stepwiseqtl* to improve the localization of QTLs and deactivated the *additive.only* option to allow for potential interactions between QTLs in the model.

We then looked into the summary output of *stepwiseqtl* to obtain information on the percentage of variance explained by each QTL, as well as the peak LOD scores, and the additive (a) and dominance (d) effects for each significant QTL, using allele N, identified in the best stepwise models for each trait. For each identified QTL, we determined the 1.5 LOD confidence interval using the *lodint* function in R/qtl. Finally, we used the *segmentsOnMap* function from QTLtools to visualize QTL segments on the genetic map, and ggplot2 (version 3.5.1; Wickham, 2011) to plot QTL effect sizes using the LOD scores obtained from the summary output of *stepwiseqtl*. To assess the distribution of estimated additive and dominance effects, we performed a separate Shapiro-Wilk normality test in R (Villasenor Alva & Estrada, 2009) for each standardized effect type using standardized phenotype residuals derived from a GLM that accounted for random effects, enabling comparisons across traits (Suppl. File 5). To identify independent QTLs-those not associated with fertility QTLs or distortion regions-we first excluded all QTLs located on chromosomes 4 and 7. Among the remaining QTLs, we removed those whose genomic intervals overlapped with Fertility Score trait QTLs on the same chromosome. Overlap was defined as any intersection between the start and end positions of the QTL intervals.

### Analysis of Whole Genome Re-sequencing Data

We used previously published resequencing data for 37 *A. nemorensis* and *A. sagittata* individuals (Dittberner *et al*., 2022). Additionally, we processed one *A. androsacea* individual used as an outgroup (Dittberner *et al*., 2022).

Reads were mapped to the new version of the *A. nemorensis* reference genome (see Genome Assembly and Annotation) using BWA mem (version 0.7.17; Li & Durbin, 2009) and default parameters. Read depth and other relevant read alignment quality control metrics were computed using QualiMap v.2.2.1 (Okonechnikov *et al*., 2016). Average read depth across all 37 samples was 23x. Variant calling was performed using GATK version 3.8 (McKenna *et al*., 2010) and duplicates were marked using PicardTools. Filtering was based on quality thresholds (DP<20; QD<2; MQ < 42; FS>60; SOR > 4; ReadPosRankSum < -6; MQRankSum < -10.5). SNPs were required to be biallelic, and sites at which more than 30% of samples were heterozygous and more than 20% missing were discarded. We polarized *A. nemorensis* and *A. sagittata* individuals with a custom Python script using *A. androsacea* as an outgroup. We performed phasing using the program shapeit2 (Delaneau *et al*., 2014), assuming a generation time of 1 year and mutation rate per generation of 7×10^-9^ and recombination rates based on the genetic map described below. From these steps, our dataset resulted in 12,868,614 SNPs before filtering, from which we extracted 4,455,806. While curating the assembly we discovered that segments on chromosomes 3, 4, 6, and 7 were inverted relative to the genetic map. Rather than remapping the reads, we lifted over the VCFs themselves: for every variant falling inside an inverted block, we mirrored its coordinate (new = start + end – old) and reverse-complemented the REF and ALT alleles. The corrected VCF files were used for all downstream analyses.

### Genome-wide Selection Scans

Using the 37 full genome samples, we identified selective sweeps using biallelic SNPs and the OmegaPlus software (Alachiotis *et al*., 2012). The OmegaPlus statistics (omega) were calculated using a grid size of 200,000 bp. We defined a minimum window size (minwin) of 50 kb and a maximum window size (maxwin) of 100 kb for computing LD values between SNPs. Outlier omega statistics, which indicated selective sweeps had occurred, were determined based on the genome-wide distribution of values. To minimize false positives arising from demographic processes, the cut-off values for the omega statistics were established using forward simulations in SLiM4 (Haller *et al.,* 2019; Haller & Messer, 2023) that were simulated under the demographic history inferred by Dittberner *et al*. (2022). In short, the demography consists of two pulses of interspecific gene flow between both species, one ancient pulse that occurred directly after the species split and one recent migration event that occurred only between the sympatric populations.

We generated 10,000 neutral datasets of 2 Mb each under the demographic history of ancient and recent migration events that occurred between the two species, assuming a fixed recombination rate for each simulated block (5.5e-8). The maximum omega value from each simulated dataset was extracted, yielding a distribution of 10,000 maximum values. The 99th percentile of this distribution that was used as the threshold to identify outlier windows indicated selective sweeps had occurred.

To optimize sweep detection, we tested multiple combinations of grid, minwin, and maxwin parameters (from 50 kb to 200 kb for grid size, and 50 kb to 100 kb for minwin and maxwin). We then applied the optimized parameters to the real dataset of *A. nemorensis* and *A. sagittata,* extracting only the sweep regions that exceeded the simulation-based threshold, which were considered high-confidence selective sweep regions.

We further investigated the potential association between selective sweeps and phenotypic traits by analyzing the overlap between the identified sweep regions and QTLs in *A. nemorensis* and *A. sagittate* (Suppl. File 7). We used only high-confidence selective sweeps in 200kb windows across the genome and overlaid these with the positions of all QTLs. We focused on the 10 percent quantile regions around the QTL peak positions to increase specificity.

### Fine-mapping the Largest QTL

To fine-map the largest QTL, which was associated with Days to Flowering trait (first QTL on chromosome 8), we identified F2 lines that were heterozygous for the QTL of interest and homozygous for other Days to Flowering QTLs; a total of 15 lines resulted (Tables S3-4). We sowed 90 F3 seeds per line (3 seeds per pot) in trays and transferred these to the cold chamber for vernalization. After two weeks, we transplanted 1 seedling into each 7x7 cm pot filled with *Topferde* (Einheitserde, Sinntal-Altengronau, Germany), resulting in 30 seedlings per line. Pots were kept in the greenhouse for six weeks, after which plants were transplanted into larger 9x9cm pots and moved to cold frames in the garden for seven weeks. During this period, leaf material was harvested from each plant, and DNA was extracted from fresh leaves using the NucleoSpin® 8 Plant II protocol. Subsequently, the plants were transplanted into new 11x11 cm pots and placed on tables in the garden under a bird-protected cage. We recorded flowering time, inflorescence height, internode length, number of shoots, plant height, stem leaf density, stem height, and rosette diameter. Two successive trials were carried out on 483 plants, the first started on 26.09.2022, the second on 16.11.2022. Plants flowered between April and June 2023.

Geneious Prime® (version 2024.0.5) was used to design multiple species-specific primers targeting the QTL region based on the *A. nemorensis* and *A. sagittata* genomes. We designed five pairs of PCR markers, which divided the QTL region into four intervals. PCR was conducted on the DNA of 407 plants to identify recombinants and locate their recombination events within the predefined intervals. We built a quantitative model using the *glm* function from the stats package (version 3.6.2) in R, accounting for both family and trial effects as fixed factors (glm(flowering_time ∼ family * (exp/tray_garden), family = quasipoisson). Then we extracted residuals and ran another model on the residuals (glm(res_ftime ∼ interval1 + interval2 + interval3 + interval4, family = gaussian)). The model was run for each interval, with the interval that best explained variation in flowering time residuals containing the flowering time QTL (Suppl. File 6). The same procedure was then applied to assess the impact of each interval on traits with overlapping QTLs in the same region as the F2 mapping population (Inflorescence Height, Rosette Diameter, and Stem Height). Parental DNA and nuclease-free water were used as positive and negative controls, respectively.

## Results

### Phenotypic Differences of Ecologically Relevant Traits

In total, 1,193 individuals germinated in the greenhouse and were set to grow in a common garden situated at the University of Cologne. All replicates of *A. nemorensis* and *A. sagittata* survived the common garden experiment (35 *A. nemorensis* and 35 *A. sagittata*). *A. nemorensis* flowered approximately 12 days earlier than A. sagittate, (*p*<0.001, Table 1). Mean values for 19 of the 22 scored traits differed between species (Table 1, Suppl. File 1, Table 1).

*A. nemorensis* individuals displayed a markedly higher number of stem leaves compared to *A. sagittata* (mean difference = 10 leaves, *p*<0.001), as expected for a trait that is often used to determine the taxonomy of these species (Titz, 1979). The rosette diameter of the parental genotypes differed significantly after 2, 3 and 4 weeks of growth (maximum *p*<0.007). Moreover, trait variance among F2 hybrids was generally larger than trait variance between their parental genotypes, suggesting transgressive segregation (Fig. 1). Although the seed production of parents did not differ significantly (Fertility Score or W.S, *p*>0.63), the low fertility observed for most F2 contrasted with the normal fertility of F1s (Fig. S1) and indicated outbreeding depression (Fig. 1).

**Figure 1.**
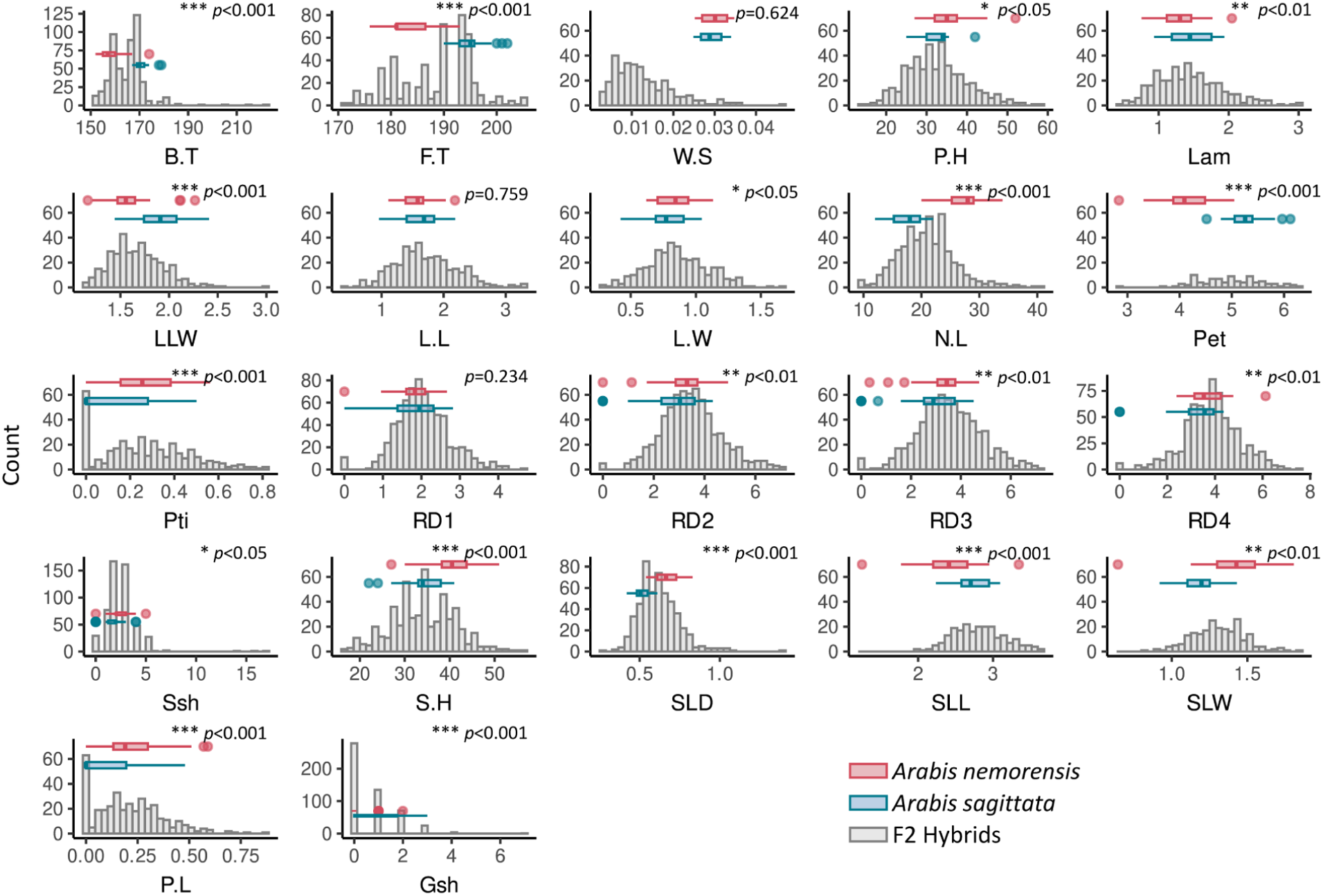
Phenotype distribution in the F2 progeny and their parental lines. Histograms illustrate the distribution of traits in the 1,193 plants of the F2 generation grown in common garden. Boxplots highlight trait variations between the two parental lines and indications of transgressive segregation in their offspring. The *p*-values and stars depict the significance of trait differences between parents * p<0.05, ** p<0.01, *** p<0.001. Abbreviations are as follows: Days to Bolting (B.T), Days to Flowering (F.T), Fertility Score - Seed Production (W.S), Inflorescence Height (P.H), Lamina Length (Lam), Lamina Length-to-Width Ratio (LLW), Leaf Length (L.L), Leaf Width (L.W), Number of Stem Leaves (N.L), Petal Length (Pet), Petiole Length (Pti), Rosette Diameter at four time points (RD1-4), Side Shoots (Ssh), Stem Height (S.H), Stem Leaf Density (SLD), Stem Leaf Length (SLL), Stem Leaf Width (SLW), Petiole Length-to-Lamina Length Ratio (P.L), and Ground Shoots (Gsh).

After four weeks of flooding, the survival rates of parental genotypes showed no significant differences (*χ^2^*(1,N=14)=0.43, *p*=0.5116).

The cross direction impacted several traits, suggesting maternal influence and potential cytoplasmic effects (Table S1). Traits such as Stem Leaf Length (SLL, *p*=3.10^-5^), Rosette Diameter at early stages (RD1, *p*=8.10^-5^), and Petiole Length (Pti, *p*=0.006) exhibited strong associations with the direction of the cross. Additionally, Petiole Length-to-Lamina Length Ratio (P.L, *p*=0.011), Rosette Diameter 2 (RD2, *p*=0.013), and Rosette Diameter 3 (RD3, *p*=0.03) also displayed significant maternal effects. *A. sagittata* maternal cytoplasmic inheritance decreased rosette lamina size on the rosette, but increased it on the stem (Table S1, Fig. S2). In contrast, the majority of traits, including Days to Bolting (B.T), Days to Flowering (F.T), Fertility Score (Seed Production or W.S), and structural traits like Inflorescence Height (P.H) and Number of Stem Leaves (N.L), showed no cytotype effects (*p*>0.05), indicating limited or negligible maternal influence.

Spearman correlation analysis of 22 traits in the F2 population revealed relationships among traits (Figs. S3-4). Traits associated with leaf shape, such as Lamina Length (Lam) and Leaf Width (L.W), exhibited strong positive correlations (r > 0.8, *p* < 0.001). Similarly, Rosette Diameter (RD1–RD4) measured at different time points showed strong positive correlations (r> 0.7, *p* < 0.001).

Developmental traits, such as Days to Bolting (B.T) and Days to Flowering (F.T), showed moderate correlations with vertical growth traits, including Stem Height (S.H) and Inflorescence Height (P.H). For instance, Inflorescence Height was positively correlated with Days to Flowering (r = 0.535, *p* < 0.001), suggesting that later-flowering individuals allocate more resources to vertical growth. In contrast, Fertility Score (Seed Production or W.S), a component of fitness, exhibited weaker and often insignificant correlations with other traits, indicating potential independence from vegetative and structural phenotypes.

Petiole traits, including Petiole Length (Pti) and its ratio to Lamina Length (P.L), which are often associated with shade avoidance strategies, were moderately correlated with rosette size and leaf shape traits such as Leaf Width (L.W). Overall, these findings highlight the interdependencies among structural traits (e.g., plant size, leaf shape), developmental traits (e.g., flowering time, rosette size over time), and fitness components in *Arabis* F2 hybrids. The statistical analyses of ecologically relevant traits in the common garden experiment provide valuable insights into the extent of phenotypic integration and the nature of resource allocation strategies in the F2 hybrid population.

### Genetic Map, Pre- and Post-zygotic Selection Distortion

To investigate the genetic architecture underlying trait variation, we used a reduced sequencing approach to determine the genotypes of 742 F2 individuals at 2,082 reliable SNP markers with the help of high-quality genome assembly (Fig. S5, Suppl. File 2, Table 1), and constructed a genetic map. The map consisted of eight linkage groups (LGs), which were numbered according to the chromosome numbers of *Arabis alpina*. The length of the linkage map was 240 cM, with 160 to 369 markers per chromosome (Table S2). Contrasting the genetic and physical distances of SNPs along chromosomes showed that recombination was higher on chromosome arms compared to in centromeric regions (Figs. S6-7).

The analysis of allele and genotype frequencies along the genome uncovered strong segregation distortions in the F2 population. A strong segregation distortion was found on chromosomes 4 and 7 with the percentage of N alleles dropping from 50% (expected) to 32% (Fig. 2A). Because the sequencing data was mapped on the high-quality *A. nemorensis* genome, we can conclude that this depletion is clearly not due to mapping biases. To disentangle prezygotic from postzygotic mechanisms of allele distortion, we estimated the difference between expected and observed genotype distributions across the genome using a chi-square test (Fig. 2B), and inferred selection coefficients against each type of homozygote. This analysis showed that the segregation distortions on chromosome 7 and on the tip of chromosome 4 were not due to postzygotic selection on the genotype. Yet, it identified 6 regions in the genome with a significant excess of heterozygotes, three of which were particularly strong on chromosomes 1 and 8. It further highlighted one region in chromosome 3 with massively depleted homozygotes.

**Figure 2.**
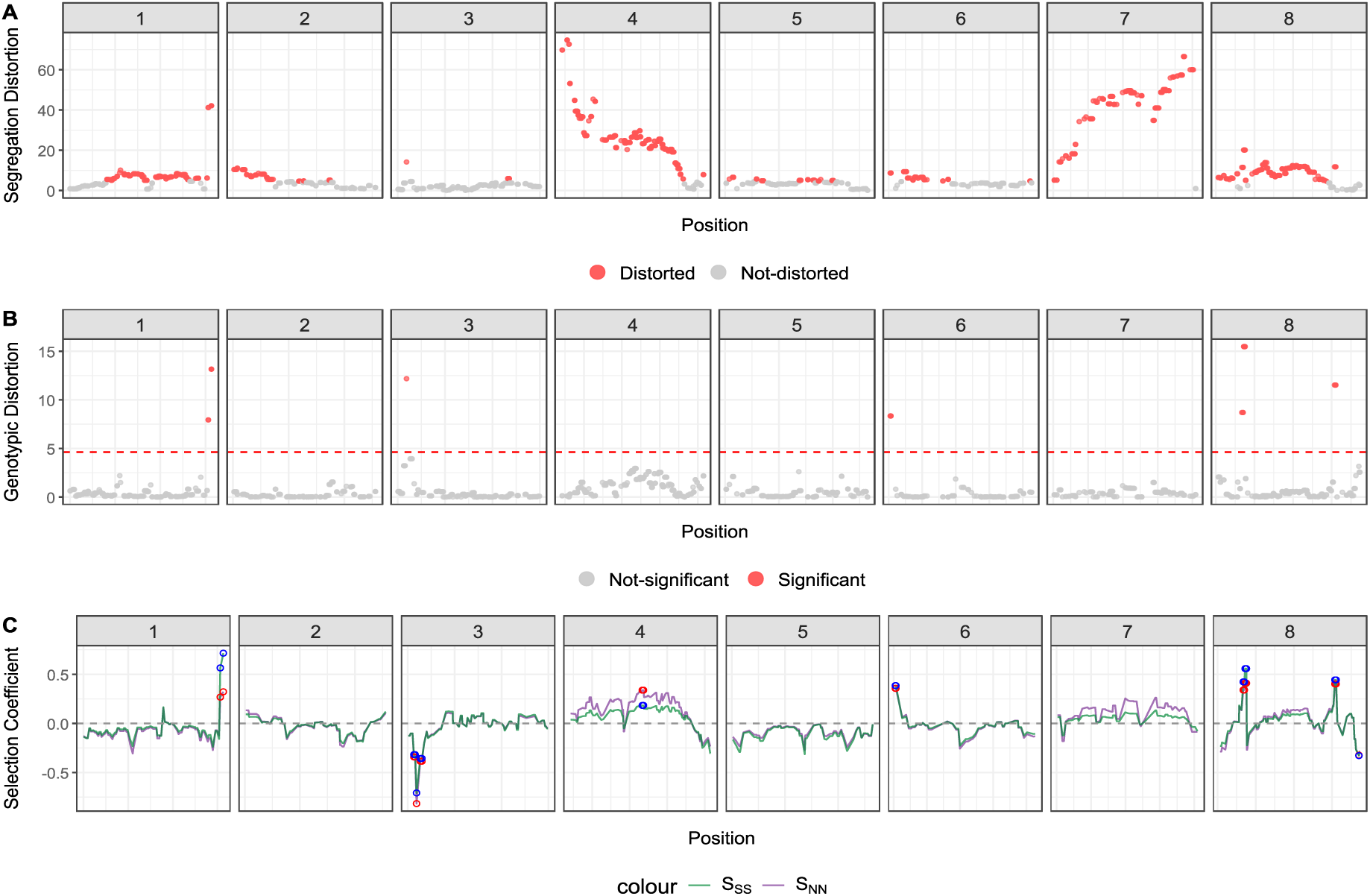
Segregation distortion, genotypic distortion and strength of selection along the genome. This figure illustrates segregation distortion, genotypic distortion, and the strength of selection across the genome in the F2 population. The x-axis represents marker positions along the genome across the eight chromosomes (2082 markers). The y-axis in (A) and (B) shows −log_10_(*p*-value); the y-axis in (C) represents the selection coefficient. (A) Gametic distortion: shown by the deviation of SNPs from expected Mendelian segregation ratios, assessed using a segregation distortion test (profileMark) with Bonferroni correction for multiple testing (threshold *p* < 2.4 × 10^−5^). Significant deviations are highlighted in red. 1,257 markers are significantly distorted. (B) Genotypic distortion: calculated as the deviation of observed genotypes from expected values according to the HWE, assessed using a chi-square test. Bonferroni-adjusted significant differences are highlighted in red. Only 47 markers surpassed the Bonferroni significance threshold. (C) Selection coefficient (*s*): calculated based on deviations from expected allele frequencies, with selection on allele S represented in green and on allele N in purple. Red and blue circles indicate markers where genotype frequencies significantly deviate from the HWE (FDR-adjusted *p* < 0.05, chi-squared test), suggesting selection at those loci. Red circles represent significant selection against the NN genotype (t); blue circles represent selection against the SS genotype (s). Loci where both t and s are significant and positive, indicate potential selection favoring the heterozygote (NS).

The extent of segregation distortion found throughout the genome appeared to be mostly due to interactions between local alleles. For example, no genetic association was observed between the distortion on chromosomes 4 and 7 (Fig. S8). Nevertheless, we find biased transmissions of parental alleles on chromosomes 2 and 5, as well as chromosomes 3 and 8 (Fig. S8).

### Genetic Architecture of Ecologically Relevant Traits

We detected significant QTLs for 20 of the 22 traits we scored (Table 2). The number of QTLs per trait ranged from 1 (Lamina Length) to 5 (Days to Flowering). The most significant QTL (LOD score > 40) was identified on chromosome 8, accounting for more than 20% of the variation in flowering time. In contrast, the lowest LOD score (3.027) was observed in one of the Leaf Width QTLs. In total, 58 QTLs were identified along the genome (Table 2). We observed overlapping QTLs for multiple traits, including 9 on chromosome 1; 3 on chromosome 2; 11 on chromosome 3; 4 on chromosome 4; 3 on chromosome 5; 2 on chromosome 6; 6 on chromosome 7; and 20 on chromosome 8 (Fig. 3). Notably, one QTL associated with the Fertility Score on chromosome 6 did not overlap with any other QTLs (Fig. 3, Table 2)

**Figure 3.**
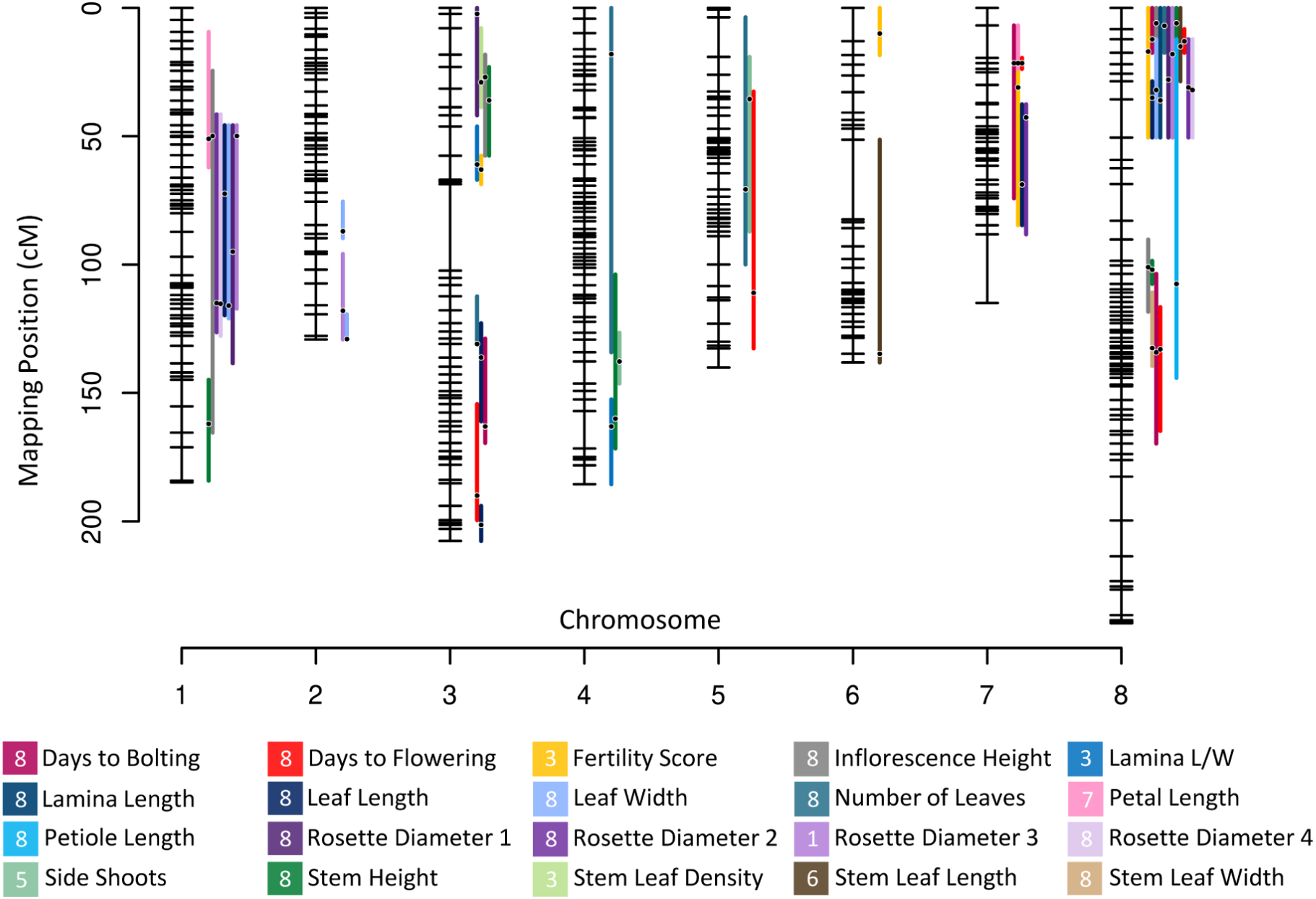
Genetic map with QTLs for ecologically relevant traits. The plot above illustrates significant QTLs that were detected and located on the genetic map. Horizontal bars represent mapped SNP markers. Gaps between bars stand for the genetic distance between SNP markers in cM (centimorgan). Traits are listed in alphabetical order. The chromosome containing the strongest QTL of each trait appears in the square.

**Table 2.** QTLs of ecologically relevant traits in the *Arabis* mapping population. This table summarizes the number, chromosomes, peak positions, LOD scores, percentages of phenotypic variance explained, and estimated additive and dominance effects of allele S for each QTL. The number of observations per trait in the F2 population is indicated. Significance differences between parental lines are denoted as follows: *** (*p* < 0.001), ** (0.001 < *p* < 0.01), * (0.01 < *p* < 0.05), . (0.05 < *p* < 0.1) (details in Suppl. File 5)

| Trait | No. of Observations | Significance of Parental Differences | Chr | Position (bp) | Position (cM) | LOD | %Variance Explained | est *a* | est *d* |
| --- | --- | --- | --- | --- | --- | --- | --- | --- | --- |
| Days to Bolting (B.T) | 631 | *** | 3 | 20040893 | 163 | 5.146 | 3.130 | 0.005914 | -0.012765 |
|  |  |  | 7 | 1835566 | 21.5 | 5.359 | 3.263 | 0.010990 | 0.002648 |
|  |  |  | 8 | 2061023 | 12.2 | 7.786 | 4.783 | -0.011989 | 0.000447 |
|  |  |  | 8 | 19500950 | 134.2 | 12.396 | 7.746 | 0.016244 | -0.003620 |
| Days to Flowering (F.T) | 578 | *** | 3 | 26882840 | 190 | 9.915 | 4.785 | 0.012242 | -0.001815 |
|  |  |  | 5 | 23418119 | 111 | 4.868 | 2.289 | -0.007092 | 0.003427 |
|  |  |  | 7 | 1835566 | 21.5 | 11.241 | 5.423 | 0.011243 | -0.001854 |
|  |  |  | 8 | 2061023 | 13 | 42.112 | 23.075 | -0.021252 | 0.008320 |
|  |  |  | 8 | 18832553 | 133 | 9.374 | 4.488 | 0.009159 | -0.005437 |
| Fertility Score (W.S) | 552 |  | 3 | 8362080 | 63 | 8.774 | 6.165 | -0.056270 | -0.366350 |
|  |  |  | 6 | 1052484 | 10 | 3.897 | 2.683 | 0.172390 | -0.055580 |
|  |  |  | 7 | 3192272 | 31 | 6.663 | 4.641 | 0.205690 | -0.053080 |
|  |  |  | 8 | 3118527 | 17 | 5.070 | 3.507 | 0.178080 | -0.028820 |
| Inflorescence Height (P.H) | 578 | . | 1 | 9004320 | 49.9 | 4.166 | 2.385 | -0.012682 | 0.062053 |
|  |  |  | 3 | 2664384 | 27 | 11.394 | 6.714 | 0.078301 | 0.016361 |
|  |  |  | 8 | 1180103 | 6 | 20.789 | 12.728 | -0.094477 | 0.076771 |
|  |  |  | 8 | 10128705 | 101 | 6.383 | 3.686 | -0.062798 | -0.038649 |
| Lamina Length (Lam) | 436 | * | 8 | 5481058 | 36.0 | 4.334 | 4.475 | -0.075590 | 0.042250 |
|  | 436 | *** | 3 | 6744303 | 61 | 15.635 | 14.890 | 0.089464 | -0.031374 |
| Lamina L/W (LLW) |  |  | 4 | 38897287 | 163 | 4.045 | 3.620 | 0.041307 | 0.028487 |
| Leaf Length (L.L) | 436 |  | 1 | 13071968 | 72.4 | 4.458 | 3.766 | 0.071867 | -0.012244 |
|  |  |  | 3 | 14255522 | 136.2 | 5.214 | 4.423 | -0.007829 | 0.118366 |
|  |  |  | 3 | 29031007 | 201.4 | 5.617 | 4.775 | 0.066584 | -0.094772 |
|  |  |  | 7 | 14342773 | 68.8 | 4.284 | 3.615 | -0.020747 | 0.086727 |
|  |  |  | 8 | 5624776 | 35 | 9.433 | 8.183 | -0.104748 | 0.036218 |
| Leaf Width (L.W) | 436 |  | 1 | 21722429 | 116 | 4.313 | 3.893 | 0.069770 | 0.045460 |
|  |  |  | 2 | 19435694 | 87 | 3.027 | 2.713 | -0.043950 | 0.056000 |
|  |  |  | 2 | 25576894 | 129 | 5.987 | 5.452 | 0.091250 | 0.012400 |
|  |  |  | 8 | 4356973 | 32 | 6.705 | 6.129 | -0.081970 | 0.036770 |
| Number of Stem Leaves (N.L) | 625 | *** | 3 | 14075550 | 130.9 | 4.581 | 2.557 | 0.025147 | 0.059693 |
|  |  |  | 4 | 7558305 | 18 | 4.001 | 2.228 | 0.056356 | 0.010451 |
|  |  |  | 5 | 16624270 | 70.7 | 5.384 | 3.014 | -0.055649 | 0.004690 |
|  |  |  | 8 | 1180103 | 7 | 28.109 | 17.139 | -0.119377 | 0.052082 |
| Petal Length (Pet) | 106 | *** | 1 | 9074627 | 51 | 4.863 | 15.820 | 0.064910 | 0.039080 |
|  |  |  | 7 | 1835566 | 21.5 | 5.689 | 18.850 | 0.079160 | 0.013480 |
| Petiole Length (Pti) | 361 | ** | 8 | 10723493 | 107.5 | 3.814 | 4.749 | -0.153200 | -0.035130 |
| Rosette Diameter 1 (RD1) | 695 |  | 1 | 17879753 | 95 | 4.498 | 2.747 | 0.068440 | -0.020590 |
|  |  |  | 3 | 623617 | 2.3 | 3.632 | 2.212 | 0.054350 | 0.023100 |
|  |  |  | 8 | 4362085 | 28 | 6.942 | 4.275 | -0.074250 | 0.032990 |
| Rosette Diameter 2 (RD2) | 700 | . | 1 | 21954996 | 115 | 6.158 | 3.703 | 0.079730 | 0.013670 |
|  |  |  | 7 | 5769578 | 42.7 | 3.839 | 2.291 | -0.056060 | 0.024000 |
|  |  |  | 8 | 4356973 | 31 | 7.822 | 4.729 | -0.085450 | 0.033910 |
|  | 696 | . | 1 | 9004320 | 49.9 | 9.389 | 5.683 | 0.102710 | -0.005208 |
| Rosette Diameter 3 (RD3) |  |  | 2 | 23541930 | 118 | 4.024 | 2.392 | 0.043216 | 0.071859 |
|  |  |  | 8 | 2718077 | 18 | 7.209 | 4.332 | -0.087105 | 0.022622 |
| Rosette Diameter 4 (RD4) | 701 | * | 1 | 21722429 | 115.3 | 4.330 | 2.678 | 0.061815 | 0.014845 |
|  |  |  | 8 | 4356973 | 32 | 7.814 | 4.888 | -0.078677 | 0.035138 |
| Side Shoots (Ssh) | 595 | * | 4 | 36501835 | 137.7 | 3.859 | 2.842 | -0.119930 | -0.038550 |
|  |  |  | 5 | 5129643 | 35.5 | 4.875 | 3.605 | 0.126580 | 0.031450 |
| Stem Height (S.H) | 625 | *** | 1 | 29847516 | 162 | 3.594 | 1.778 | -0.029247 | 0.036282 |
|  |  |  | 3 | 4035375 | 36 | 13.598 | 6.984 | 0.074366 | 0.005981 |
|  |  |  | 4 | 38897287 | 160 | 4.386 | 2.177 | 0.042699 | -0.004189 |
|  |  |  | 8 | 1180103 | 6 | 26.851 | 14.500 | -0.091994 | 0.070533 |
|  |  |  | 8 | 10506892 | 102 | 7.858 | 3.951 | -0.059677 | -0.004408 |
| Stem Leaf Density (SLD) | 624 | *** | 3 | 3491752 | 29 | 5.760 | 4.162 | -0.052426 | -0.018081 |
| Stem Leaf Length (SLL) | 259 | ** | 6 | 21070306 | 134.8 | 5.918 | 9.356 | 0.046349 | 0.011155 |
|  |  |  | 8 | 3118527 | 15 | 3.967 | 6.163 | -0.035380 | 0.011240 |
| Stem Leaf Width (SLW) | 259 | *** | 8 | 18832576 | 132.5 | 13.228 | 20.958 | -0.084603 | 0.017725 |

The strongest QTLs for traits such as Days to Bolting, Days to Flowering, Inflorescence Height, Leaf Length and Width, Number of Stem Leaves, and Rosette Diameter after 1, 2, and 4 weeks, as well as Stem Height and Stem Leaf Width, were all located on chromosome 8. Additionally, strongest QTLs for Petal Length, Rosette Diameter 3, Side Shoots, Stem Leaf Density, and Stem Leaf Length were identified on chromosomes 7, 1, 5, 3, and 6, respectively. For all traits, the additive effect of the *A. nemorensis* allele was either positive or negative, confirming that transgressive genetic variation can arise in most traits in this selfing species by fixing a new combination of alleles (Table 2).

The distribution of fertility scores, which was measured as the weight of seeds contained in 10 siliques, indicated that F2 individuals tended to be less fertile than the parental lineages. The genetic architecture of fertility score variation was dominated by a large-effect QTL (LOD = 8.774) on chromosome 3 (Fig. S9), which explained more than 30% of the phenotypic variation and involved inter-allelic incompatibility at position 8,362,080 bp on chromosome 3 (QTL interval: 6,744,303 to 9,601,991 bp, Fig. S9). Individuals with the NS heterozygous genotype at this marker displayed markedly lower fertility. In addition, three smaller QTLs explaining 2.683%, 4.641%, and 3.507% of the variation were found on chromosomes 6, 7, and 8, respectively. It can be concluded that the genetic basis of outbreeding depression is relatively simple in this population.

Of the 22 traits scored, Ground Shoots (Gsh), Petiole Length-to-Lamina Length ratio (P.L), and Survival to Flooding revealed no significant QTL. For two of these three traits (Ground Shoots, Petiole Length-to-Lamina Length ratio), significant differences between species were observed. The absence of QTLs for these traits suggests they may have a polygenic genetic basis, and the effect of individual variants may be too small to be detected.

We conclude that the two parental lineages differ genetically in many traits, with size effects of various sizes. The number of detected QTLs allows us to describe the size effects of genetic variation that can be generated by hybridization in this system. The standardized additive effects were normally distributed (Shapiro-Wilk: W = 0.9744, *p* = 0.2573, Fig. 4), ranging from a minimum of -0.15 to a maximum of 0.20 with a median of 0.008 (mean = 0.005, s.d.= 0.08, skewness = 0.29, kurtosis = 2.6). QTLs with the largest effect controlled flowering time, stem height, inflorescence height, leaf number and lamina shape contributing to the heavy tail of the distribution of additive effect (Figs. 4, S10). Standardized dominance effects instead approximated a logistic distribution (Shapiro-Wilk: W = 0.683, *p* < 0.001), ranging from -0.36 to 0.11 with a median of 0.012 (mean = 0.008, s.d. = 0.062, skewness =-3.70, kurtosis = 23.5). The dominance effect of the fertility score QTL on chromosome 3 was an outlier to this distribution (Fig. 4).

**Figure 4.**
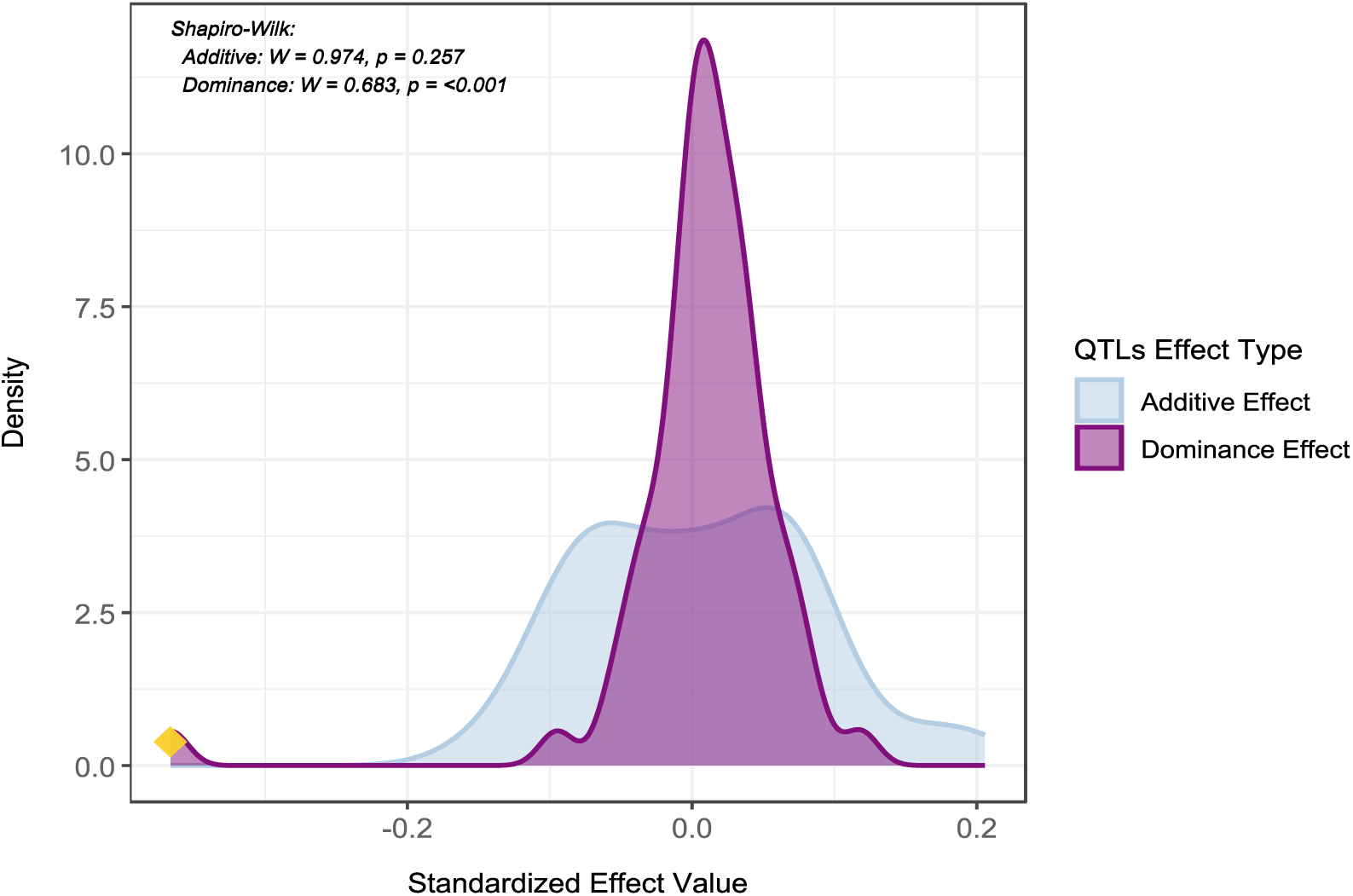
Distribution of standardized QTL estimated additive and dominance effects. The figure above displays the distribution of standardized estimated additive and dominance effects for QTLs associated with ecologically relevant traits identified in the F2 population. Each curve represents the density of standardized effect sizes for a distinct type of genetic effect: additive effects (in light blue) and dominance effects (in purple). The yellow diamond marks the dominance effect of the strongest QTL for Fertility Score (W.S) on chromosome 3. Effect values were estimated using interval mapping on standardized phenotype residuals derived from a GLM that accounted for random effects, enabling comparisons across traits. Density distributions illustrate the range and frequency of QTL contributions. Additive effects showed a Gaussian distribution (Shapiro-Wilk: W = 0.974, *p* = 0.257), whereas dominance effects approximated a logistic distribution after excluding the outlier dominance effect of a QTL on chromosome 3 (Shapiro-Wilk: W = 0.683, *p* < 0.001).

### Genes Responsible for Flowering Time

The largest QTL detected in this study impacts the timing of flowering and is located on chromosome 8, where two well-known genes that control flowering time are found: *FLC*, a regulator involved in the vernalization pathway, and *CONSTANS*, a regulator involved in the photoperiod pathway (Andrés & Coupland, 2012). *FLC* has been shown to be a key variant shaping flowering time in many Brassicaceae species, including the congeneric species *Arabis alpina* (Albani *et al*., 2012). The gene *CONSTANS* is also located within the boundaries of this QTL. In order to test whether one of these candidate genes was responsible for the variation, we selected 15 F2 individuals that were heterozygous in the QTL region of chromosome 8 and homozygous on the other QTLs, and we grew 30 of their seeds. A total of 410 plants were assessed for flowering time in 15 F3 families and two trials (Fig. S11, Tables S3-4).

With 410 plants and a QTL region that was ∼17 cM, we expected 80 recombinants. Interestingly, we identified 138 recombinants, suggesting that the recombination rate had been slightly underestimated in the F2 population. We compared four models to identify the chromosomal fragments that best explained variation in flowering time, after accounting for all other factors of the experimental design. Using Akaike’s criterion and comparing *p*-values, we identified fragment 2 as the most likely to contain the variants causing a difference in flowering time (Fig. 5, *p*=0.0118). This segment ranges from position 1,831,324 bp to position 2,125,083 bp at the beginning of chromosome 8. Although this ∼300 kb region does not contain *FLC* or *CO*, it contains 64 other genes, 31 of which have a known ortholog in *Arabidopsis thaliana*. Only 1 of these, *TFL1*, is known to regulate flowering time in *A. thaliana* and *A. alpina*, with a loss of function inducing early flowering in both species (Cerise *et al*., 2023). *A. sagittata TFL1* differs from the *A. nemorensis* copy by 2 amino acid exchanges: Asparagine 3 is changed to Isoleucine and Serine 50 is changed Tyrosine. The *A. nemorensis* version at both sites is conserved in most Brassicaceae, like *A. thaliana*, where *TFL1* function has been experimentally verified (Andrés & Coupland, 2012). Nevertheless, both positions are not strictly conserved.

**Figure 5.**
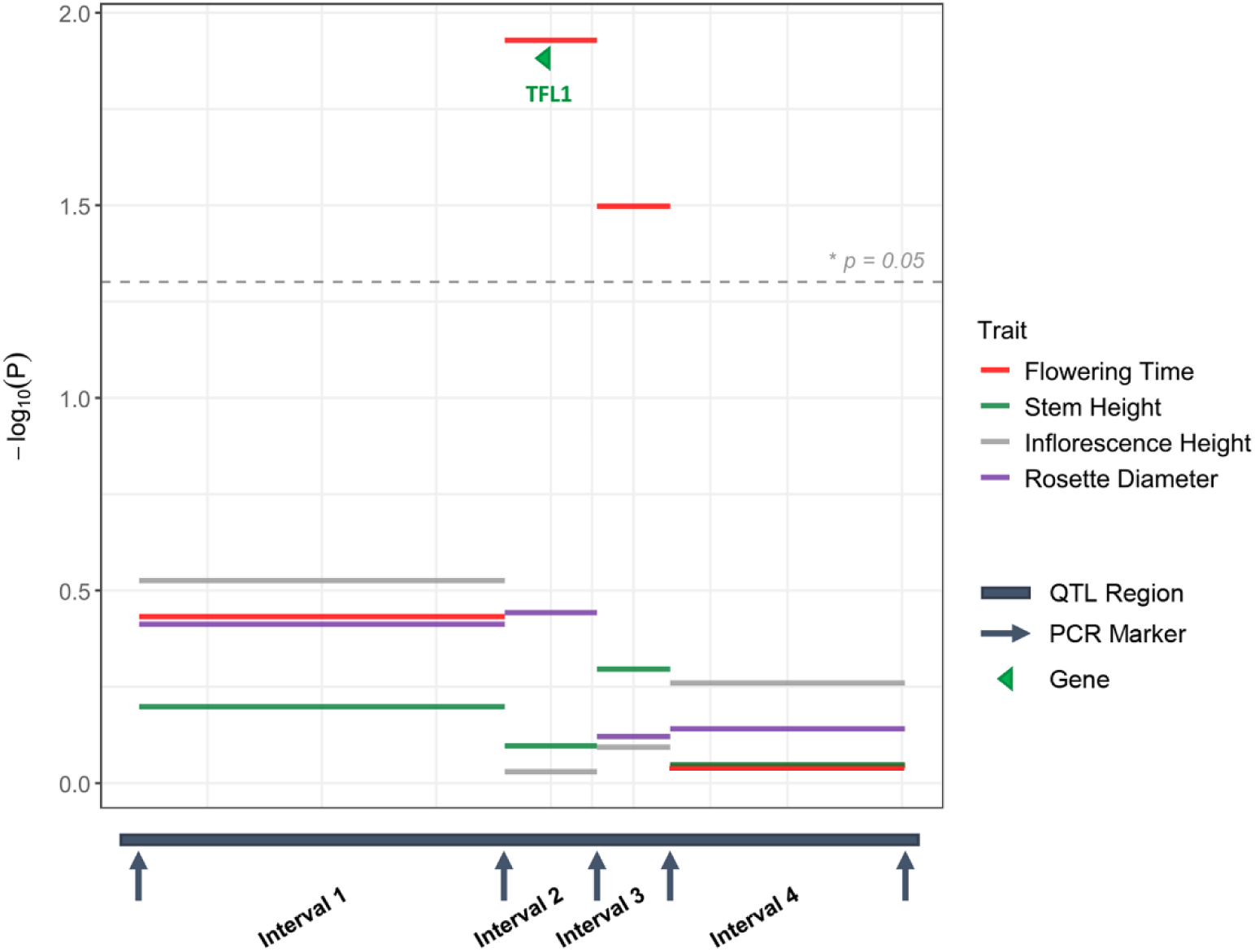
Association between 4 DNA fragments and flowering time within 15 F3 families segregated for the chr8 flowering time QTL. The figure illustrates the role of genomic intervals within the QTL region in explaining variations in Flowering Time, Inflorescence Height, Rosette Diameter, and Stem Height. The x-axis represents region of the strongest Flowering Time QTL, divided into intervals defined by species-specific primers with adjusted lengths for fine mapping. The dashed line indicates the significance threshold. Different bar colors represent different traits. The *TFL1* gene was detected in interval 2, which shows the strongest effect on flowering time.

In contrast to most *TFL1* genes from the Brassicales, several species within (*Capsella rubella*, *Sinapis alba*) and outside of this group are missing the start codon and protein translation starts at the conserved Methionine codon at position 4. At position 50, Serine is the most common amino acid not only in the Brassicales group, but also in other plant groups, but other amino acids (Alanine, Threonine, Phenylalanine) are found in other Brassicaceae *TFL*1 sequences. Because *A. sagittata* carries derived amino acids at two moderately conserved positions, gene activity may be decreased compared to the more ancestral *A. nemorensis* allele. Fine-mapping, however, allowed us to exclude the role of flowering loci such as *FLC* or *CO* that locate on the same chromosome arm of chromosome 8.

Although there were QTLs for Inflorescence Height, Rosette Diameter, and Stem Height, on chromosome 8 in F2s, the chromosomal region that was fine mapped did not explain the variation of these traits in F3s. Inflorescence Height, Rosette Diameter and Stem Height are thus controlled by QTL(s) independent of the QTL for Flowering Time in the *TFL1*-containing fragment, with the S allele advancing flowering compared to the N allele.

### Overlap of Parental QTLs with the History of Selection in the Parental Lineages

In order to test whether genetic differences between lineages were influenced by selection for ecological specialization in the two parental lineages, prior to hybridization, we used previously published population genomics data to determine regions with signature of selective sweeps (Dittberner *et al*., 2022). We detected 35 and 82 genome regions carrying signatures of selective sweeps in *A. nemorensis* and *A.sagittata*, respectively, which prompted us to ask whether variable traits were disproportionately targeted by natural selection in their lineage or origin (Fig. S12). Many of the 58 QTL regions spanned large parts of chromosomes, sometimes overlapping with multiple independent selective sweeps (Fig. S13). We tested whether the distance between QTL peak and selective sweep was shorter than expected by a permutation test. QTL peaks did not tend to be located close to selective sweep signatures. We also did not detect significant overlaps between regions with distorted segregation and regions carrying selective sweeps (not shown). The genetic variation segregating in the F2 population thus does not appear to reflect the history of selection in the hybridizing lineages, yet some individual QTLs may reflect recent selection. For example, sweeps overlapped with 5 of the 7 genomic regions with QTLs for rosette size (Fig. S12). In contrast, no signature of a putative selective sweep was found in the *TFL1* region.

## Discussion

Hybridization between species is pervasive in both past and contemporary ecosystems (Lewontin & Birch, 1966; Stull *et al*., 2023). By bringing together alleles that contribute to different ecological specialization, it can facilitate the emergence of a new allelic combination that may surpass the performance of the parental genotypes and be fixed via selfing. Hybridization may therefore become pivotal in the rescue of endangered selfing species with very low levels of genetic diversity (Frankham, 2015). Here, we take advantage of a hybridization hotspot we identified previously to disentangle the genomic and ecological properties of genetic variation released after hybridization, some of these properties may propagate in the population via selfing. An F2 generation was obtained by crossing individuals representative of the two species in the sympatric population where they naturally hybridize (Dittberner *et al*., 2019; Dittberner *et al*., 2022).

The genetic analysis of allelic transmission in the F2 population first revealed the genomic barriers that skew allele segregation in hybrid offspring. Indeed, several regions of the genome displayed highly distorted transmission. For the two most strongly distorted regions on chromosomes 4 and 7, the genotypic composition of the F2 fits Mendelian expectations, suggesting that the distortion happens before fertilization, as a result of biased gamete formation. Both of these distortions drive the *A. sagittata* allele to high frequencies in the F2 offspring population. The predominance of one allele among mature gametes produced in a heterozygous individual represents a very strong evolutionary force called the meiotic drive (Sandler & Novitski, 1957; Pinkas *et al*., 1985; Clarke *et al*., 2004; Domínguez *et al*., 2005). The meiotic drive can massively accelerate the fixation of alleles linked to the driver locus. Driver alleles may outcompete non-driver alleles during pollen tube growth in heterozygotic selfing plants, thereby biasing the paternal transmission of alleles in favor of the offspring (e.g., Snow & Spira, 1996; Aronen *et al*., 2002; Lankinen *et al*., 2009). In female gametes, any molecular change that favors allele transmission into the polar body would lead to preferential maternal transmission (Fishman & Kelly, 2015; Finseth *et al*., 2015). Here, since genotype frequencies fulfil the HWE, the driver alleles should outcompete the non-driver alleles in both male and female gametogenesis (Malik, 2005). Many mechanisms can lead to gametic drive in plants, because gametes undergo haploid cell division, a step that does not exist in animals (Finseth, 2023). As a consequence, about 60% of plant genes are expressed in the haploid phase (Chettoor *et al*., 2014; Rutley & Twell, 2015; Klepikova *et al*., 2016), and variants have been identified in genes controlling cell division in gametes of *A. thaliana* (Parker *et al*., 2025).

In addition to segregation distortion at the gametic level, we also detected regions in the genome that experienced biased transmission as a result of the distorted frequency of heterozygotes. One locus on chromosome 3, for example, was markedly depleted in heterozygotes, despite equal transmission of the two alleles. This locus was close, but distinct from a second locus that decreased the seed production of heterozygotes by 30%. This pattern, which is also known as allelic underdominance, characterizes the genome of a pair of alleles in *A. thaliana* (Smith *et al*., 2011). The causal variant mapped to structural changes in a tandem array of duplicated genes and altered kinase activity (Smith *et al*., 2011). In selfing species and their hybrids, underdominance will not pose a significant threat to the equal transmission of alleles provided that F1 hybrids are viable – as is the case in this study system– because half of the descendants of heterozygote individuals will return to homozygosity.

Allelic underdominance is a form of Dobzhansky-Muller (DM) incompatibility (Dobzhansky, 1936; Muller, 1942). DM incompatibilities arise after alleles are fixed in the diverging parental lineages without being naturally selected to function together. Interactions between alleles segregating at different loci can also cause DM incompatibilities, but, interestingly, no segregation distortions were detected in any major interlocus in this system, nor did any interactive QTLs determine fertility. DM incompatibilities have been studied extensively in many taxa (Bomblies & Weigel, 2007; Masly & Presgraves, 2007; White *et al*., 2011; Schumer *et al*., 2014; Zuellig & Sweigart, 2018; Coughlan & Matute, 2020). Although such incompatibilities are assumed to evolve completely neutrally, they can also result from local adaptation (Alcázar *et al*., 2014).

However, whereas only 2 loci showed underdominance, 6 loci showed selection for heterozygote genotypes. This discrepancy indicates that in this system the compensation of deleterious variants in heterozygote regions is more predominant than the emergence of DM incompatibilities (Clo *et al*., 2021). Indeed, 6 regions in the genome show an excess of heterozygotes. This imbalance indicates that the low effective population sizes of the two species, as well as their relatively recent origin, will have increased the fixation of deleterious alleles faster than they will have fixed new incompatible mutations (Simons & Sella, 2016; Dittberner *et al*., 2022). As recombination and selfing will proceed in further generations after hybridization, high-fitness individuals are expected to arise that will have purged these variants. The interplay between purging and recombination has been shown to cause heterogeneous rates of introgression along the genome of hybridizing taxa (Schumer *et al*., 2018).

The pattern of past introgression in the *Arabis* system is in fact heterogeneous along the genome (Dittberner *et al*., 2022). Yet, our work shows that the selective removal of deleterious variants is probably not the only force affecting this system today. The massive transmission advantage of the *A. sagittata* alleles on chromosomes 4 and 7 implies that any allele that is located close to the driver of the distortion will readily introgress into the local *A. nemorensis* population, whether it is deleterious or ecologically relevant. Linked QTLs of small effect in the vicinity of the distortion, however, will be hard to detect. Other approaches, such as, for example, transcriptome analyses, are needed to determine the potential ecological relevance of genetic variation hitchhiking with gametic driver alleles.

The genomics of allele transmission is clearly shaping the consequences of hybridization in this system. Yet, segregation distortions, incompatibilities, and overdominance were detectable on a limited number of loci. Due to their simple genetic basis, the loci are unlikely to block genetic exchanges (Li *et al*., 2022). The genetic architecture of the comprehensive panel of traits quantifying growth rate, growth form, the transition to flowering, plant height and the production of seeds allowed us to examine the extent to which gene flow at ecologically relevant QTLs was hindered by genomic barriers in this system. Results demonstrated that the parental lineages differ genetically in most traits, with about 48% of the QTLs contributing to this variation, that is, most traits are independent of the loci affecting segregation or fertility in the F2 generations (Table S5).

Hybrid F2 offspring often exhibited phenotypic values beyond the range of the parental lineages, particularly for plant height, rosette diameter, leaf length and flowering time. Indeed, although the two parental genotypes did not differ in the timing of flowering, it was for this trait that the largest QTL was detected on chromosome 8. Our study showed that changes in flowering time in *A. nemorensis* and *A. sagittata* hybrids are not linked to the well-known genes controlling flowering time, *FLC* or *CO* (reviewed in Alonso-Blanco *et al*., 2009; Andrés & Coupland, 2012). Instead, the fine-mapping of the largest QTL defined a genetic region that contains only one candidate gene for flowering time. This gene, first described as *TFL1* in *A. thaliana,* accelerates flowering by decreasing the number of vegetative side buds in this species (Alvarez *et al*., 1992; Moraes *et al*., 2019). In the perennial species *A. alpina*, it further interacts with the age pathway to prevent juvenile plants from responding to vernalization, thereby controlling the transition out of the juvenile stage, an additional developmental transition of central importance for perennials (Wang & Albani *et al*., 2011). Because this flowering time variant is independent from all segregation distortions and fertility alleles, it appears as a potential adaptive trait likely to be reshaped as a result of gene flow. Studies of flowering time provided some of the most remarkable examples of contemporary adaptation to climate change in plants (Hancock *et al*., 2024 submitted), and adaptive introgression of flowering time alleles have been documented in several cases (Le Corre *et al*., 2002; Todesco *et al*., 2020; Wang *et al*., 2021). Environmental responses to global climate change include advancing flowering time: many plants induce earlier flowering as a strategy to escape warmer temperatures (e.g., Anderson *et al*., 2012; Cai *et al*., 2019; Siegmund *et al*., 2016; Tun *et al*., 2021).

Which additional, genetically different traits may be adaptive in natural communities remains to be determined. Larger leaves, greater lateral spread and lower specific leaf area, all of which are variable in the descendants of hybrid individuals in this system, have been associated with competitive advantage in the ephemeral wetland of vernal pool plant communities (Kraft *et al*., 2015). Interestingly, the *A. nemorensis* maternal background increased rosette diameter, especially at early stages, while the *A. sagittata* maternal background increased stem leaf length. Both of these traits can improve light capture, but their relative fitness relevance should depend on the trade-off between survival and competition (Lundgren & Des Marais, 2020). Backcrossing of *A. nemorensis* pollen on *A. sagittata* mother plants appeared to occur more frequently among hybrids (Dittberner *et al*., 2019; Dittberner *et al*., 2022), which would thus promote an increase in stem leaf length. Therefore, genetic and maternal variants may contribute to the emergence of a particularly competitive genotype in the dense plant communities of floodplain meadows. In this context, it is intriguing that signatures of putative selective sweeps in the parental species were found for many of the QTLs determining variation among rosette size. Recent selection in the parental lineages however, has not been systematically associated with variable QTLs, so that much of the variation manifested in the F2 generation forms a new base of variants that has not yet been shaped by natural selection.

As the climate changes and land-use intensifies, the native floodplain habitat is increasingly exposed to drought events as well as flooding, both of which will affect plant survival in natural environments and neither of which could be quantified in this common garden experiment (Colloff *et al*., 2016). In situ analyses of the performance of hybrid offspring are therefore needed to shed light on the genetic basis of variation in survival rates. We conclude that there is sufficient genetic diversity in our system for hybridization to generate novelty. The prospects of hybridization between these species, however, will depend on how selective pressures act on genetically variable traits within the architecture of allelic transmission.

## Acknowledgments

This work was supported by the Deutsche Forschungsgemeinschaft (DFG [ME2742/13-1, TRR341 and EXC 2048/1–390686111). We thank the Cologne Center for Genomics (CCG), the Regionales Rechenzentrum der Universität zu Köln (RRZK), the German Network for Bioinformatics Infrastructure (de.NBI) and ELIXIR-DE (Forschungszentrum Jülich and W-de.NBI-001, W-de.NBI-004, W-de.NBI-008, W-de.NBI-010, W-de.NBI-013, W-de.NBI-014, W-de.NBI-016, W-de.NBI-022). We thank K. Schneeberger, B. Hüttel, A. Hancock, M. Stetter, I. Calic and M. Hajheidari for valuable feedback. This work benefited from the contribution of Y. Özoglan to data collection.

## Author contributions

N.R. designed and performed the RADseq and QTL mapping analyses, conducted flowering-time experiments and analyses, and wrote the manuscript. L.M. contributed to the whole-genome sequencing (WGS) analyses and wrote the WGS section of the manuscript. L.H. assisted with RADseq data analysis. K.K. assisted with genotype correction. A.S.K. generated the genome annotations. T.A., C.I.D., S.A., R.Y.W, and N.R. generated the genome assemblies. G.S. and B. S. contributed to data interpretation. A.T. and J.d.M. conceived and supervised the project, contributed to data interpretation, and reviewed and edited the manuscript.

## Data Availability

FASTQ files of the F2 population, genome assemblies, and annotations are available at the European Nucleotide Archive (ENA) under project number [PRJEB89863]. All supplementary figures can be found at Supplementary Figures. All scripts, VCF file, and phenotypic datasets used in this study are available on GitHub via the following links:

- Suppl. File 1. Phenotypic analyses: https://github.com/nedarahnama/Contemporary_hybridization/tree/master/01_phenotypic_analyses
- Suppl. File 2. Genome assembly: https://github.com/nedarahnama/Contemporary_hybridization/tree/master/02_genome_assembly
- Suppl. File 3. RADseq analysis: https://github.com/nedarahnama/Contemporary_hybridization/tree/master/03_rad_seq
- Suppl. File 4. Genetic map construction: https://github.com/nedarahnama/Contemporary_hybridization/tree/master/04_genetic_map
- Suppl. File 5. QTL mapping analysis: https://github.com/nedarahnama/Contemporary_hybridization/tree/master/05_qtl_mapping
- Suppl. File 6. Flowering time fine-mapping analysis: https://github.com/nedarahnama/Contemporary_hybridization/tree/master/06_fine_mapping
- Suppl. File 7. Sweep detection: https://github.com/Luker121/OverlapSweepQTL/tree/main

## References

1. Abbott RJ. 2017. Plant speciation across environmental gradients and the occurrence and nature of hybrid zones. Journal of Systematics and Evolution 55.

2. Abbott R, Albach D, Ansell S, Arntzen JW, Baird SJE, Bierne N, Boughman J, Brelsford A, Buerkle CA, Buggs R, et al. 2013. Hybridization and speciation. Journal of Evolutionary Biology 26.

3. Alachiotis N, Stamatakis A, Pavlidis P. 2012. OmegaPlus: A scalable tool for rapid detection of selective sweeps in whole-genome datasets. Bioinformatics 28.

4. Albani MC, Castaings L, Wötzel S, Mateos JL, Wunder J, Wang R, Reymond M, Coupland G. 2012. PEP1 of *Arabis alpina* Is Encoded by Two Overlapping Genes That Contribute to Natural Genetic Variation in Perennial Flowering. PLoS Genetics 8.

5. Alcázar R, von Reth M, Bautor J, Chae E, Weigel D, Koornneef M, Parker JE. 2014. Analysis of a Plant Complex Resistance Gene Locus Underlying Immune-Related Hybrid Incompatibility and Its Occurrence in Nature. PLoS Genetics 10.

6. Alonge M, Lebeigle L, Kirsche M, Jenike K, Ou S, Aganezov S, Wang X, Lippman ZB, Schatz MC, Soyk S. 2022. Automated assembly scaffolding using RagTag elevates a new tomato system for high-throughput genome editing. Genome Biology 23.

7. Alonso-Blanco C, Aarts MGM, Bentsink L, Keurentjes JJB, Reymond M, Vreugdenhil D, Koornneef M. 2009. What has natural variation taught us about plant development, physiology, and adaptation? Plant Cell 21.

8. Alvarez J, Guli CL, Yu X -H, Smyth DR. 1992. terminal flower: a gene affecting inflorescence development in *Arabidopsis thaliana*. The Plant Journal 2.

9. Anderson JT, Panetta AM, Mitchell-Olds T. 2012. Evolutionary and Ecological Responses to Anthropogenic Climate Change. Plant Physiology 160: 1728–1740.

10. Anderson E, Stebbins GL. 1954. HYBRIDIZATION AS AN EVOLUTIONARY STIMULUS. Evolution 8.

11. Andrés F, Coupland G. 2012. The genetic basis of flowering responses to seasonal cues. Nature Reviews Genetics 13.

12. Andrews S. 2010. FastQC: a quality control tool for high throughput sequence data. 2010. Https://Www.Bioinformatics.Babraham.Ac.Uk/Projects/Fastqc/.

13. Aronen T, Nikkanen T, Harju A, Tiimonen H, Häggman H. 2002. Pollen competition and seed-siring success in *Picea abies*. Theoretical and Applied Genetics 104.

14. Becker M, Gruenheit N, Steel M, Voelckel C, Deusch O, Heenan PB, McLenachan PA, Kardailsky O, Leigh JW, Lockhart PJ. 2013. Hybridization may facilitate in situ survival of endemic species through periods of climate change. Nature Climate Change 3.

15. Blanckaert A, Sriram V, Bank C. 2023. In search of the Goldilocks zone for hybrid speciation II: hard times for hybrid speciation? Evolution 77.

16. Bomblies K, Weigel D. 2007. Hybrid necrosis: Autoimmunity as a potential gene-flow barrier in plant species. Nature Reviews Genetics 8.

17. Bontrager M, Angert AL. 2019. Gene flow improves fitness at a range edge under climate change. Evolution Letters 3.

18. Brauer CJ, Sandoval-Castillo J, Gates K, Hammer MP, Unmack PJ, Bernatchez L, Beheregaray LB. 2023. Natural hybridization reduces vulnerability to climate change. Nature Climate Change 13.

19. Broman KW, Wu H, Sen Ś, Churchill GA. 2003. R/qtl: QTL mapping in experimental crosses. Bioinformatics 19.

20. Buerkle CA, Morris RJ, Asmussen MA, Rieseberg LH. 2000. The likelihood of homoploid hybrid speciation. Heredity 84.

21. Burmeier S, Eckstein RL, Donath TW, Otte A. 2011. Plant pattern development during early post-restoration succession in grasslands-a case study of *Arabis nemorensis*. Restoration Ecology 19.

22. Cai Z, Zhou L, Ren NN, Xu X, Liu R, Huang L, Zheng XM, Meng QL, Du YS, Wang MX, et al. 2019. Parallel Speciation of Wild Rice Associated with Habitat Shifts. Molecular Biology and Evolution 36.

23. Carey K, Ganders FR. 1980. Heterozygote Advantage at the Fruit Wing Locus in *Plectritis congesta* (Valerianaceae). Evolution 34.

24. Catchen J, Hohenlohe PA, Bassham S, Amores A, Cresko WA. 2013. Stacks: An analysis tool set for population genomics. Molecular Ecology 22.

25. Ceballos G, Ehrlich PR, Raven PH. 2020. Vertebrates on the brink as indicators of biological annihilation and the sixth mass extinction. Proceedings of the National Academy of Sciences 117: 13596–13602.

26. Cerise M, da Silveira Falavigna V, Rodríguez-Maroto G, Signol A, Severing E, Gao H, van Driel A, Vincent C, Wilkens S, Iacobini FR, et al. 2023. Two modes of gene regulation by *TFL1* mediate its dual function in flowering time and shoot determinacy of *Arabidopsis*. Development (Cambridge) 150.

27. Cheng H, Concepcion GT, Feng X, Zhang H, Li H. 2021. Haplotype-resolved de novo assembly using phased assembly graphs with hifiasm. Nature Methods 18.

28. Cheng H, Jarvis ED, Fedrigo O, Koepfli KP, Urban L, Gemmell NJ, Li H. 2022. Haplotype-resolved assembly of diploid genomes without parental data. Nature Biotechnology.

29. Chettoor AM, Givan SA, Cole RA, Coker CT, Unger-Wallace E, Vejlupkova Z, Vollbrecht E, Fowler JE, Evans MMS. 2014. Discovery of novel transcripts and gametophytic functions via RNA-seq analysis of maize gametophytic transcriptomes. Genome Biology 15.

30. Chunco AJ. 2014. Hybridization in a warmer world. Ecology and Evolution 4.

31. Clarke HJ, Khan TN, Siddique KHM. 2004. Pollen selection for chilling tolerance at hybridisation leads to improved chickpea cultivars. Euphytica 139.

32. Clo J, Ronfort J, Gay L. 2021. Fitness consequences of hybridization in a predominantly selfing species: insights into the role of dominance and epistatic incompatibilities. Heredity 127.

33. Cochrane SKJ, Andersen JH, Berg T, Blanchet H, Borja A, Carstensen J, Elliott M, Hummel H, Niquil N, Renaud PE. 2016. What is marine biodiversity? Towards common concepts and their implications for assessing biodiversity status. Frontiers in Marine Science 3.

34. Colloff MJ, Lavorel S, Wise RM, Dunlop M, Overton IC, Williams KJ. 2016. Adaptation services of floodplains and wetlands under transformational climate change. Ecological Applications 26.

35. Cooper BS, Sedghifar A, Nash WT, Comeault AA, Matute DR. 2018. A Maladaptive Combination of Traits Contributes to the Maintenance of a *Drosophila* Hybrid Zone. Current Biology 28.

36. [Core Writing Team HL and JR (eds. ). 2022. Climate Change 2023: Synthesis Report. Contribution of Working Groups I, II and III to the Sixth Assessment Report of the Intergovernmental Panel on Climate Change. IPCC 13.

37. Le Corre V, Roux F, Reboud X. 2002. DNA polymorphism at the *FRIGIDA* gene in *Arabidopsis thaliana*: Extensive nonsynonymous variation is consistent with local selection for flowering time. Molecular Biology and Evolution 19.

38. Coughlan JM, Matute DR. 2020. The importance of intrinsic postzygotic barriers throughout the speciation process: Intrinsic barriers throughout speciation. Philosophical Transactions of the Royal Society B: Biological Sciences 375.

39. Cowie RH, Bouchet P, Fontaine B. 2022. The Sixth Mass Extinction: fact, fiction or speculation? Biological Reviews 97.

40. Danecek P, Auton A, Abecasis G, Albers CA, Banks E, DePristo MA, Handsaker RE, Lunter G, Marth GT, Sherry ST, et al. 2011. The variant call format and VCFtools. Bioinformatics 27.

41. Delaneau O, Marchini J, McVeanh GA, Donnelly P, Lunter G, Myers S, Gupta-Hinch A, Iqbal Z, Mathieson I, Rimmer A, et al. 2014. Integrating sequence and array data to create an improved 1000 Genomes Project haplotype reference panel. Nature Communications 5.

42. Delaneau O, Ongen H, Brown AA, Fort A, Panousis NI, Dermitzakis ET. 2017. A complete tool set for molecular QTL discovery and analysis. Nature Communications 8.

43. Dittberner H, Becker C, Jiao WB, Schneeberger K, Hölzel N, Tellier A, de Meaux J. 2019. Strengths and potential pitfalls of hay transfer for ecological restoration revealed by RAD-seq analysis in floodplain *Arabis* species. Molecular Ecology 28.

44. Dittberner H, Tellier A, De Meaux J. 2022. Approximate Bayesian Computation Untangles Signatures of Contemporary and Historical Hybridization between Two Endangered Species. Molecular Biology and Evolution 39.

45. Dobzhansky T. 1936. STUDIES ON HYBRID STERILITY. II. LOCALIZATION OF STERILITY FACTORS IN DROSOPHILA PSEUDOOBSCURA HYBRIDS. Genetics 21.

46. Domínguez E, Cuartero J, Fernández-Muñoz R. 2005. Breeding tomato for pollen tolerance to low temperatures by gametophytic selection. Euphytica 142.

47. Durand NC, Robinson JT, Shamim MS, Machol I, Mesirov JP, Lander ES, Aiden EL. 2016a. Cell Systems 3: 99–101.

48. Durand NC, Shamim MS, Machol I, Rao SS, Huntley MH, Lander ES, Aiden EL. 2016b. Cell Systems 3: 95–98.

49. Eichenberg D, Bowler DE, Bonn A, Bruelheide H, Grescho V, Harter D, Jandt U, May R, Winter M, Jansen F. 2021. Widespread decline in Central European plant diversity across six decades. Global Change Biology 27.

50. Ellstrand NC, Schierenbeck KA. 2000. Hybridization as a stimulus for the evolution of invasiveness in plants? Proceedings of the National Academy of Sciences of the United States of America 97.

51. Finseth F. 2023. Female meiotic drive in plants: mechanisms and dynamics. Current Opinion in Genetics and Development 82.

52. Finseth FR, Dong Y, Saunders A, Fishman L. 2015. Duplication and adaptive evolution of a key centromeric protein in mimulus, a genus with female meiotic drive. Molecular Biology and Evolution 32.

53. Fishman L, Kelly JK. 2015. Centromere-associated meiotic drive and female fitness variation in *Mimulus*. Evolution 69.

54. Frankham R. 2015. Genetic rescue of small inbred populations: meta-analysis reveals large and consistent benefits of gene flow. Molecular Ecology 24.

55. Goulet BE, Roda F, Hopkins R. 2017. Hybridization in plants: Old ideas, new techniques[OPEN]. Plant Physiology 173.

56. Haley CS, Knott SA. 1992. A simple regression method for mapping quantitative trait loci in line crosses using flanking markers. Heredity 69.

57. Haller BC, Galloway J, Kelleher J, Messer PW, Ralph PL. 2019. Tree-sequence recording in SLiM opens new horizons for forward-time simulation of whole genomes. Molecular Ecology Resources 19.

58. Haller BC, Messer PW. 2023. SLiM 4: Multispecies Eco-Evolutionary Modeling. American Naturalist 201.

59. Hancock A, Portalier S, Fulgione A, Stetter MG, de Meaux J. 2024. The molecular basis of adaptation to climatic factors and range change in plants. Annual reviews in Ecology and Evolution. Submitted.

60. Hansen MM, Olivieri I, Waller DM, Nielsen EE. 2012. Monitoring adaptive genetic responses to environmental change. Molecular Ecology 21.

61. Hölzel N. 2005. Seedling recruitment in flood-meadow species: The effects of gaps, litter and vegetation matrix. Applied Vegetation Science 8.

62. Hooper JNA, Kennedy JA, Quinn RJ. 2002. Biodiversity ‘hotspots’, patterns of richness and endemism, and taxonomic affinities of tropical Australian sponges (Porifera). Biodiversity and Conservation 11.

63. Hopkins R. 2013. Reinforcement in plants. New Phytologist 197.

64. Karl R, Koch MA. 2014. Phylogenetic signatures of adaptation: The *Arabis hirsuta* species aggregate (Brassicaceae) revisited. Perspectives in Plant Ecology, Evolution and Systematics 16.

65. Klepikova A V., Kasianov AS, Gerasimov ES, Logacheva MD, Penin AA. 2016. A high resolution map of the *Arabidopsis thaliana* developmental transcriptome based on RNA-seq profiling. Plant Journal 88.

66. Kraft TS, Wright SJ, Turner I, Lucas PW, Oufiero CE, Supardi Noor MN, Sun IF, Dominy NJ. 2015. Seed size and the evolution of leaf defences. Journal of Ecology 103.

67. Lankinen S, Maad J, Armbruster WS. 2009. Pollen-tube growth rates in *Collinsia heterophylla* (Plantaginaceae): One-donor crosses reveal heritability but no effect on sporophytic-offspring fitness. Annals of Botany 103.

68. Lewontin RC, Birch LC. 1966. Hybridization as a Source of Variation for Adaptation to New Environments. Evolution 20.

69. Li H, Durbin R. 2009. Fast and accurate short read alignment with Burrows-Wheeler transform. Bioinformatics 25.

70. Li H, Handsaker B, Wysoker A, Fennell T, Ruan J, Homer N, Marth G, Abecasis G, Durbin R. 2009. The Sequence Alignment/Map format and SAMtools. Bioinformatics 25.

71. Li J, Schumer M, Bank C. 2022. Imbalanced segregation of recombinant haplotypes in hybrid populations reveals inter- and intrachromosomal Dobzhansky-Muller incompatibilities. PLoS Genetics 18.

72. Lundgren MR, Des Marais DL. 2020. Life History Variation as a Model for Understanding Trade-Offs in Plant–Environment Interactions. Current Biology 30.

73. Ma Y, Marczewski T, Xue D, Wu Z, Liao R, Sun W, Marczewski J. 2019. Conservation implications of asymmetric introgression and reproductive barriers in a rare primrose species. BMC Plant Biology 19.

74. Malik HS. 2005. *Mimulus* finds centromeres in the driver’s seat. Trends in Ecology and Evolution 20.

75. Mallet J. 2007. Hybrid speciation. Nature 446: 279–283.

76. Marçais G, Kingsford C. 2011. A fast, lock-free approach for efficient parallel counting of occurrences of k-mers. Bioinformatics 27: 764–770.

77. Martin M. 2011. Cutadapt removes adapter sequences from high-throughput sequencing reads. EMBnet.journal 17.

78. Masly JP, Presgraves DC. 2007. High-resolution genome-wide dissection of the two rules of speciation in *Drosophila*. PLoS Biology 5.

79. Mathar W, Kleinebecker T, Hölzel N. 2015. Environmental variation as a key process of co-existence in flood-meadows. Journal of Vegetation Science 26.

80. McKenna A, Hanna M, Banks E, Sivachenko A, Cibulskis K, Kernytsky A, Garimella K, Altshuler D, Gabriel S, Daly M, et al. 2010. The genome analysis toolkit: A MapReduce framework for analyzing next-generation DNA sequencing data. Genome Research 20.

81. Moore JS, Hendry AP. 2009. Can gene flow have negative demographic consequences? Mixed evidence from stream threespine stickleback. Philosophical Transactions of the Royal Society B: Biological Sciences 364.

82. Moraes TS, Dornelas MC, Martinelli AP. 2019. *FT*/*TFL1*: Calibrating plant architecture. Frontiers in Plant Science 10.

83. Moran BM, Payne CY, Powell DL, Iverson ENK, Donny AE, Banerjee SM, Langdon QK, Gunn TR, Rodriguez-Soto RA, Madero A, et al. 2024. A lethal mitonuclear incompatibility in complex I of natural hybrids. Nature 626.

84. Muller HJ. 1942. Isolating mechanisms, evolution and temperature. Biol. Symp 6.

85. Nocchi G, Wang J, Yang L, Ding J, Gao Y, Buggs RJA, Wang N. 2023. Genomic signals of local adaptation and hybridization in Asian white birch. Molecular Ecology 32.

86. Okonechnikov K, Conesa A, García-Alcalde F. 2016. Qualimap 2: Advanced multi-sample quality control for high-throughput sequencing data. Bioinformatics 32.

87. Parker MT, Amar S, Campoy JA, Krause K, Tusso S, Marek M, Huettel B, Schneeberger K. 2025. Scalable eQTL mapping using single-nucleus RNA-sequencing of recombined gametes from a small number of individuals. PLOS Biology 23.

88. Peñalba J V., Runemark A, Meier JI, Singh P, Wogan GOU, Sánchez-Guillén R, Mallet J, Rometsch SJ, Menon M, Seehausen O, et al. 2024. The Role of Hybridization in Species Formation and Persistence. Cold Spring Harbor Perspectives in Biology.

89. Pfennig KS, Kelly AL, Pierce AA. 2016. Hybridization as a facilitator of species range expansion. Proceedings of the Royal Society B: Biological Sciences 283.

90. Pinkas R, Zamir D, Ladizinsky G. 1985. Allozyme divergence and evolution in the genusLens. Plant Systematics and Evolution 151: 131–140.

91. Powell DL, García-Olazábal M, Keegan M, Reilly P, Du K, Díaz-Loyo AP, Banerjee S, Blakkan D, Reich D, Andolfatto P, et al. 2020. Natural hybridization reveals incompatible alleles that cause melanoma in swordtail fish. Science 368.

92. Presgraves DC. 2010. The molecular evolutionary basis of species formation. Nature Reviews Genetics 11.

93. Ranallo-Benavidez TR, Jaron KS, Schatz MC. 2020. GenomeScope 2.0 and Smudgeplot for reference-free profiling of polyploid genomes. Nature Communications 11.

94. Rhymer JM, Simberloff D. 1996. Extinction by hybridization and introgression. Annual Review of Ecology and Systematics 27.

95. Rieseberg LH, Raymond O, Rosenthal DM, Lai Z, Livingstone K, Nakazato T, Durphy JL, Schwarzbach AE, Donovan LA, Lexer C. 2003. Major ecological transitions in wild sunflowers facilitated by hybridization. Science 301.

96. Rivera-Colón AG, Catchen J. 2022. Population Genomics Analysis with RAD, Reprised: Stacks 2. In: Methods in Molecular Biology.

97. Rosser N, Seixas F, Queste LM, Cama B, Mori-Pezo R, Kryvokhyzha D, Nelson M, Waite-Hudson R, Goringe M, Costa M, et al. 2024. Hybrid speciation driven by multilocus introgression of ecological traits. Nature 628: 811–817.

98. Rutley N, Twell D. 2015. A decade of pollen transcriptomics. Plant Reproduction 28.

99. Sandler L, Novitski E. 1957. Meiotic Drive as an Evolutionary Force. The American Naturalist 91.

100. Schlaepfer MA, Lawler JJ. 2023. Conserving biodiversity in the face of rapid climate change requires a shift in priorities. Wiley Interdisciplinary Reviews: Climate Change 14.

101. Schnittler M, Günther KF. 1999. Central European vascular plants requiring priority conservation measures - An analysis from national Red Lists and distribution maps. Biodiversity and Conservation 8.

102. Schumer M, Brandvain Y. 2016. Determining epistatic selection in admixed populations. Molecular ecology 25.

103. Schumer M, Cui R, Powell DL, Dresner R, Rosenthal GG, Andolfatto P. 2014. High-resolution mapping reveals hundreds of genetic incompatibilities in hybridizing fish species. eLife 2014.

104. Schumer M, Cui R, Rosenthal GG, Andolfatto P. 2015. Reproductive Isolation of Hybrid Populations Driven by Genetic Incompatibilities. PLoS Genetics 11.

105. Schumer M, Xu C, Powell DL, Durvasula A, Skov L, Holland C, Blazier JC, Sankararaman S, Andolfatto P, Rosenthal GG, et al. 2018. Natural selection interacts with recombination to shape the evolution of hybrid genomes. Science 360.

106. Seehausen O. 2004. Hybridization and adaptive radiation. Trends in Ecology and Evolution 19.

107. Siegmund JF, Wiedermann M, Donges JF, Donner R V. 2016. Impact of temperature and precipitation extremes on the flowering dates of four German wildlife shrub species. Biogeosciences 13.

108. Sim SB, Corpuz RL, Simmonds TJ, Geib SM. 2022. HiFiAdapterFilt, a memory efficient read processing pipeline, prevents occurrence of adapter sequence in PacBio HiFi reads and their negative impacts on genome assembly. BMC Genomics 23.

109. Simons YB, Sella G. 2016. The impact of recent population history on the deleterious mutation load in humans and close evolutionary relatives. Current Opinion in Genetics and Development 41.

110. Smit AFA, Hubley R, Grenn P. 2015. RepeatMasker Open-4.0. RepeatMasker Open-4.0.7.

111. Smith LM, Bomblies K, Weigel D. 2011. Complex evolutionary events at a tandem cluster of *Arabidopsis thaliana* genes resulting in a single-locus genetic incompatibility. PLoS Genetics 7.

112. Snow AA, Spira TP. 1996. Pollen-tube competition and male fitness in *Hibiscus moscheutos*. Evolution 50.

113. Staudinger MD, Grimm NB, Staudt A, Carter SF, Chapin FS, Kareiva P, Ruckelshaus M, Stein BA. 2012. Impacts of climate change on biodiversity, ecosystems, and ecosystem services: technical input to the 2013 National Climate Assessment. Cooperative Report to the 2013 National Climate Assessment.

114. Stull GW, Pham KK, Soltis PS, Soltis DE. 2023. Deep reticulation: the long legacy of hybridization in vascular plant evolution. Plant Journal 114.

115. Taylor J, Butler D. 2017. R package ASMap: Efficient genetic linkage map construction and diagnosis. Journal of Statistical Software 79.

116. Theissinger K, Fernandes C, Formenti G, Bista I, Berg PR, Bleidorn C, Bombarely A, Crottini A, Gallo GR, Godoy JA, et al. 2023. How genomics can help biodiversity conservation. Trends in Genetics 39.

117. Titz W. 1979. Die Interfertilitätsbeziehungen europäischer Sippen der *Arabis hirsuta*-Gruppe (Brassicaceae). Plant Systematics and Evolution 131.

118. Todesco M, Owens GL, Bercovich N, Légaré JS, Soudi S, Burge DO, Huang K, Ostevik KL, Drummond EBM, Imerovski I, et al. 2020. Massive haplotypes underlie ecotypic differentiation in sunflowers. Nature 584.

119. Todesco M, Pascual MA, Owens GL, Ostevik KL, Moyers BT, Hübner S, Heredia SM, Hahn MA, Caseys C, Bock DG, et al. 2016. Hybridization and extinction. Evolutionary Applications 9.

120. Tun W, Yoon J, Jeon JS, An G. 2021. Influence of Climate Change on Flowering Time. Journal of Plant Biology 64.

121. Villasenor Alva JA, Estrada EG. 2009. A generalization of Shapiro-Wilk’s test for multivariate normality. Communications in Statistics - Theory and Methods 38.

122. Wang R, Albani MC, Vincent C, Bergonzi S, Luan M, Bai Y, Kiefer C, Castillo R, Coupland G. 2011. Aa tfl1 confers an age-dependent response to vernalization in perennial *Arabis alpina*. Plant Cell 23.

123. Wang Z, Jiang Y, Bi H, Lu Z, Ma Y, Yang X, Chen N, Tian B, Liu B, Mao X, et al. 2021. Hybrid speciation via inheritance of alternate alleles of parental isolating genes. Molecular Plant 14.

124. Wang RJ, White MA, Payseur BA. 2015. The Pace of hybrid incompatibility evolution in house mice. Genetics 201.

125. White MA, Steffy B, Wiltshire T, Payseur BA. 2011. Genetic dissection of a key reproductive barrier between nascent species of house mice. Genetics 189.

126. White MA, Stubbings M, Dumont BL, Payseur BA. 2012. Genetics and evolution of hybrid male sterility in house mice. Genetics 191.

127. Wickham H. 2011. ggplot2. Wiley Interdisciplinary Reviews: Computational Statistics 3: 180–185.

128. Zuellig MP, Sweigart AL. 2018. Gene duplicates cause hybrid lethality between sympatric species of *Mimulus*. PLoS Genetics 14.

