## Supplementary material for "Contemporary hybridization among *Arabis* floodplain species creates opportunities for adaptation": Suppl. data

### Supplementary Figures

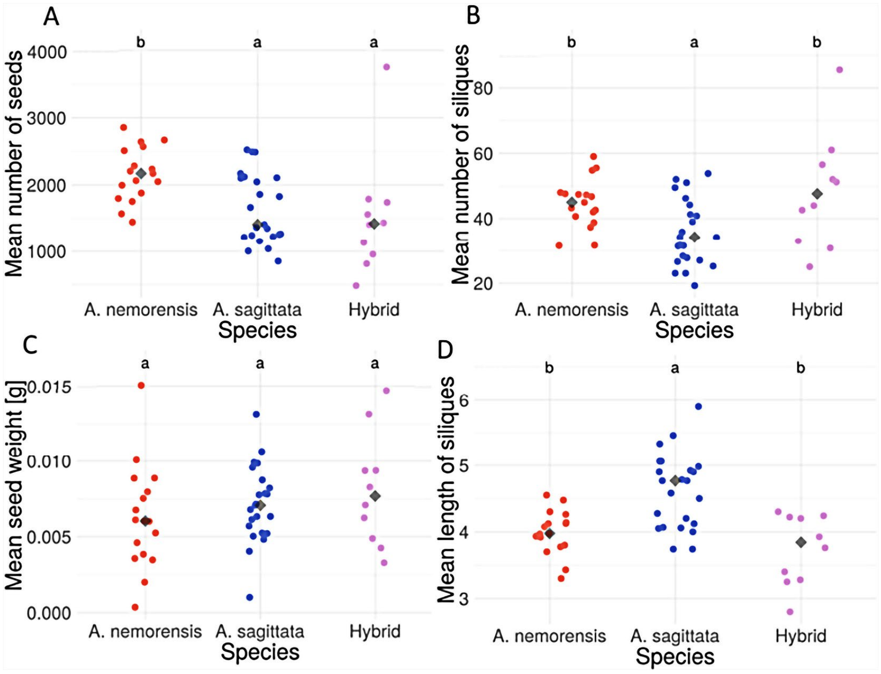


**Figure S1. Phenotypic variation in fitness-related traits in parental species and F1 hybrids.** Each point represents the mean value per accession from three populations (described in Dittberner *et al.* 2022), including number of seeds, number of siliques, mean seed weight, silique length; diamonds indicate the median. Letters above data points denote significant differences between groups ($p$<0.05). The parental species (*A. nemorensis* and *A. sagittata*) were reciprocally crossed to produce F1 hybrids. The fitness of F1 was comparable to that of the parental species. Seedlings were grown in the greenhouse of the Experimental Garden of the University of Cologne, and the seeds of the first generation of selfing (F2) were collected.

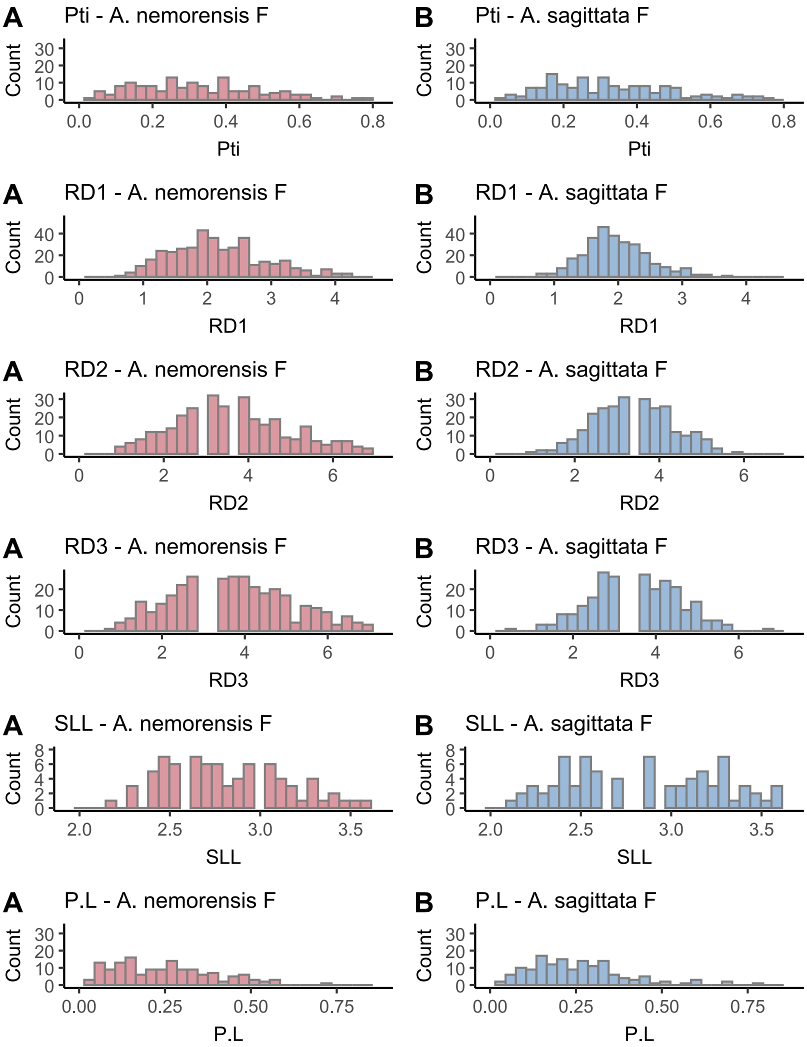


**Figure S2. Phenotype distribution in the F2 progeny and the effect of cross-direction.** Histograms show the distribution of phenotypic traits in 1,193 F2 individuals grown in a common garden, highlighting traits for which the direction of the cross has a significant effect. Colors indicate the maternal species: red represents crosses in which *A. nemorensis* is the female parent (A), and blue represents crosses in which *A. sagittata* is the female parent (B). The *p*-values for the effect of *A. sagittata* as the female parent on each trait are as follows: Pti: 0.00591, RD1: 0.00008, RD2: 0.01358, RD3: 0.03091, SLL: 0.00003, P.L: 0.01190. When *A. sagittata* is the female parent, petiole length (Pti), stem leaf length (SLL), and the petiole-to-lamina length ratio (P.L) are significantly increased. In contrast, rosette diameter at the first three time points (RD1–RD3) is significantly decreased. For more details, see Table S1.

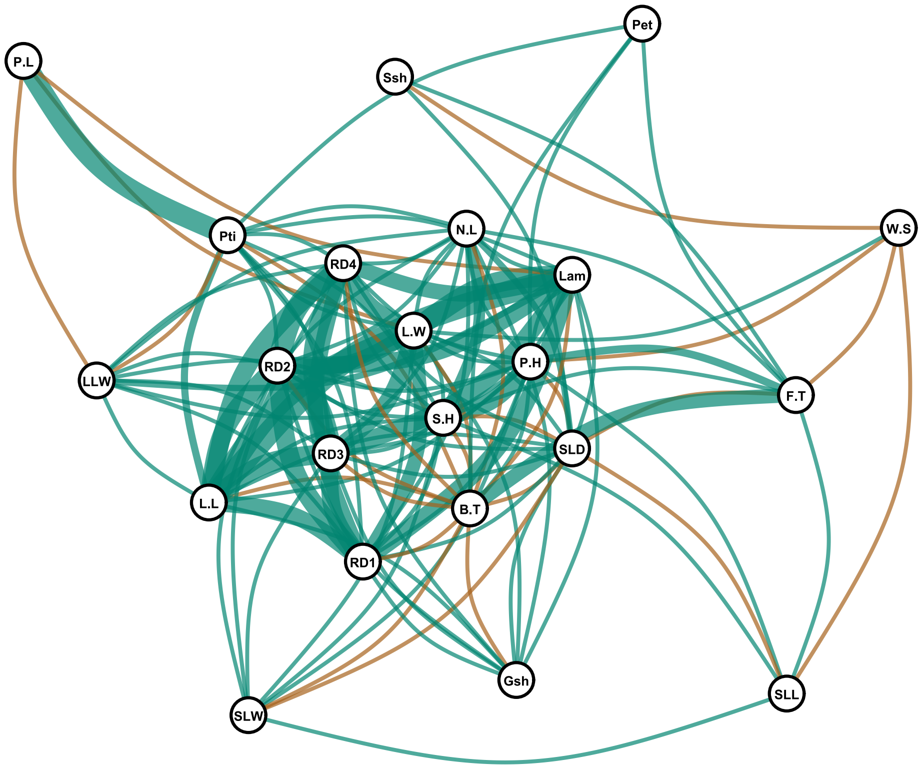


**Figure S3. Correlation network of phenotypic traits.** The figure above illustrates the relationships among phenotypic traits measured in the common garden experiment. Spearman's rank correlation coefficients were conducted in R to quantify pairwise relationships between traits, with residuals from quasi-Poisson models accounting for variation due to experimental blocks and cross-direction. Pairwise correlations were calculated with pairwise deletions for missing data. Edges represent statistically significant correlations ($\alpha = 0.05$), based on raw *p*-values. The thickness of each edge reflects the strength of the correlation, scaled non-linearly to emphasize stronger relationships: correlations with $|r| \leq 0.5$ are drawn with thinner, uniform widths, whereas those with $|r| > 0.5$ increase in thickness up to a maximum. Partially transparent curved edges improve visual clarity. Positive correlations are shown in green and negative in brown, using a colorblind-friendly palette. The visualization was generated using the ggraph and igraph packages in R, the measured phenotypes include: Days to Bolting (B.T), Days to Flowering (F.T), Fertility Score - Seed Production (W.S), Inflorescence Height (P.H), Lamina Length (Lam), Lamina Length-to-Width Ratio (LLW), Leaf Length (L.L), Leaf Width (L.W), Number of Stem Leaves (N.L), Petal Length (Pet), Petiole Length (Pti), Rosette Diameter at four time points (RD1-4), Side Shoots (Ssh), Stem Height (S.H), Stem Leaf Density (SLD), Stem Leaf Length (SLL), Stem Leaf Width (SLW), Petiole Length-to-Lamina Length Ratio (P.L), and Ground Shoots (Gsh).

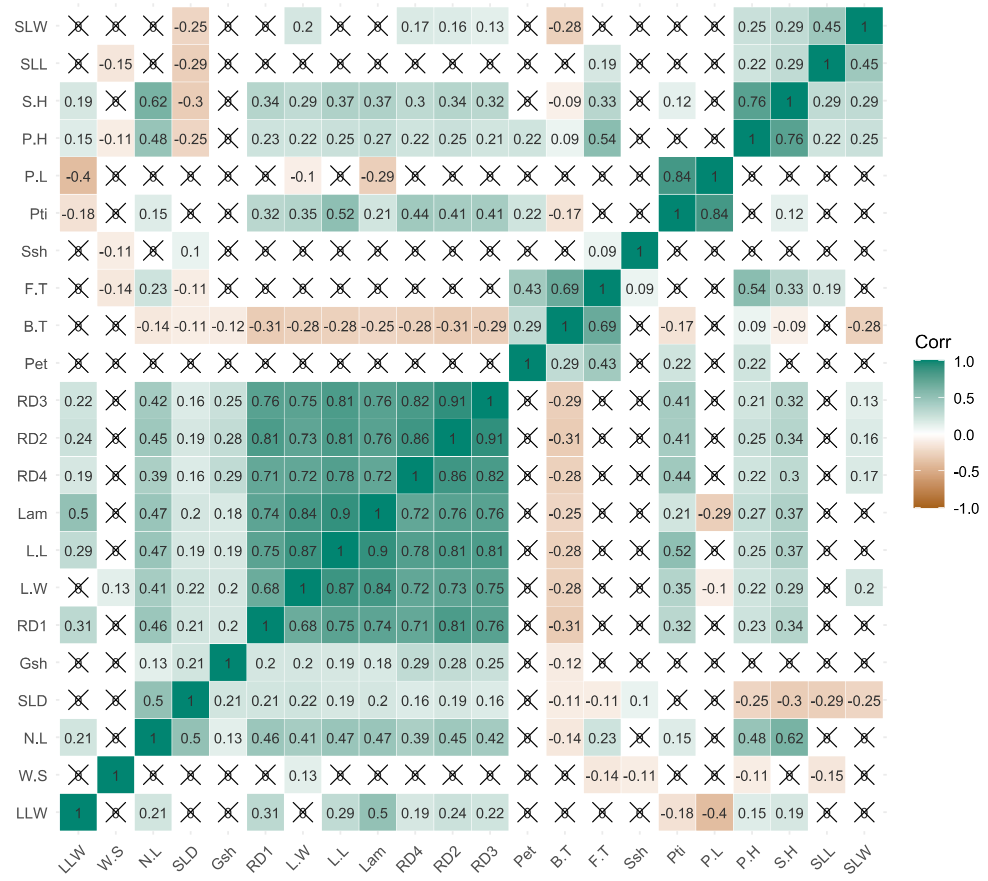


**Figure S4. Correlation heatmap of phenotypic traits.** This heatmap provides a complementary visualization of the full correlation matrix using Spearman's rank correlation coefficients. The heatmap is organized hierarchically and displays significant correlations with corresponding labels. A color gradient was applied to distinguish correlation strengths. Positive correlations are shown in green and negative in brown, using a colorblind-friendly palette. Non-significant correlations ($\alpha > 0.05$) are marked with a cross (×) on their corresponding cells, allowing them to be quickly distinguished from statistically supported relationships. Both visualizations (Figs. S3 and S4) provided complementary insights into the complex association patterns among traits, helping to interpret phenotypic relationships in F2 plants within the common garden experiment. The phenotypes we measured were aas follows: Days to Bolting (B.T), Days to Flowering (F.T), Fertility Score - Seed Production (W.S), Inflorescence Height (P.H), Lamina Length (Lam), Lamina Length-to-Width Ratio (LLW), Leaf Length (L.L), Leaf Width (L.W), Number of Stem Leaves (N.L), Petal Length (Pet), Petiole Length (Pti), Rosette Diameter at four time points (RD1-4), Side Shoots (Ssh), Stem Height (S.H), Stem Leaf Density (SLD), Stem Leaf Length (SLL), Stem Leaf Width (SLW), Petiole Length-to-Lamina Length Ratio (P.L), and Ground Shoots (Gsh).

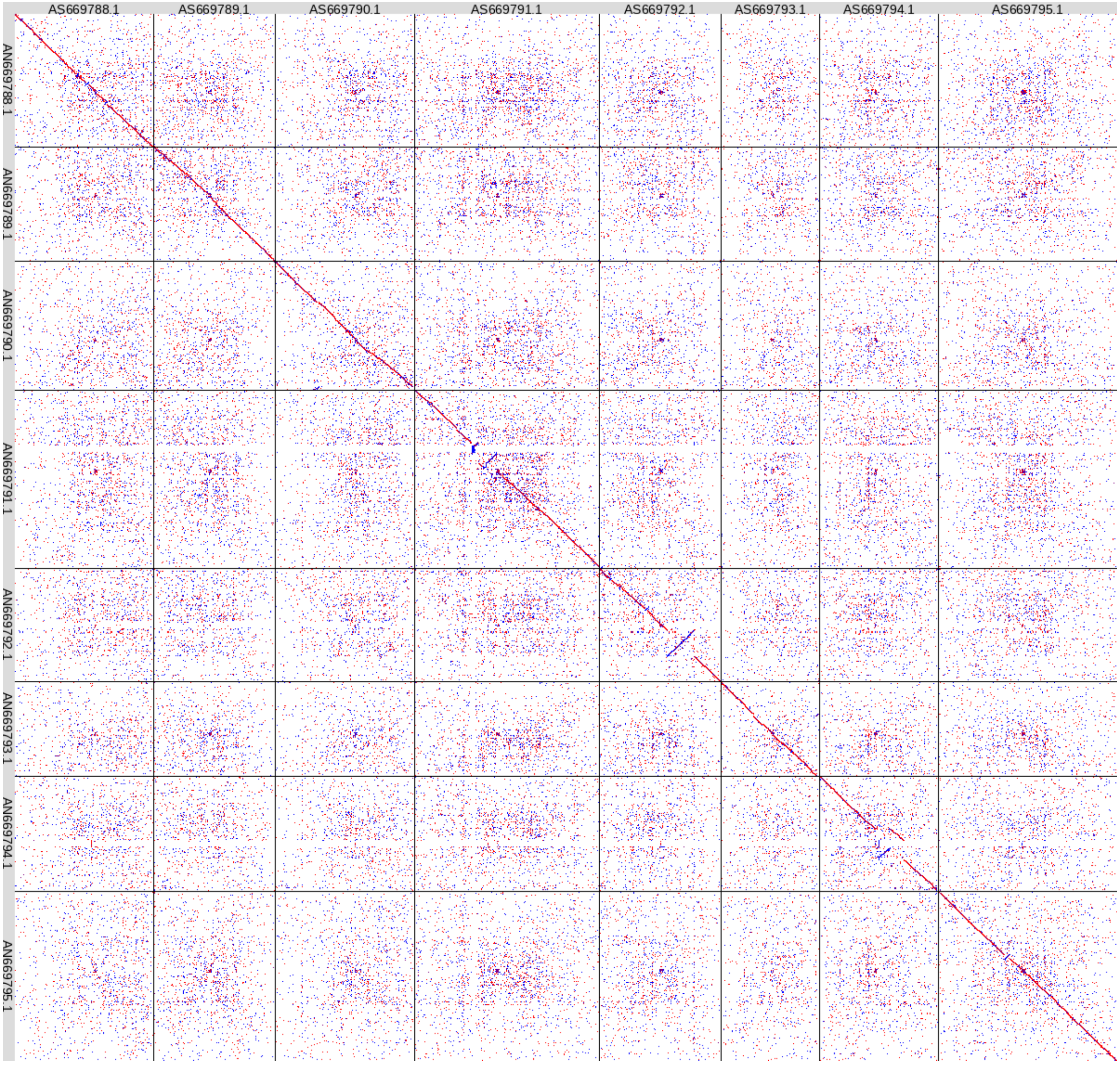


**Figure S5. Synteny and rearrangement plot between *A. nemorensis* and *A. sagittata* genomes.** The dot plot illustrates synteny and the localization of genomic rearrangements between the final RagTag-generated assemblies of *A. nemorensis* and *A. sagittata*. The x-axis represents scaffolds (1 to 8) of *A. sagittata* (from left to right); the y-axis represents scaffolds (1 to 8) of *A. nemorensis* (from top to bottom). Potential inversions are observed on chromosomes 3, 4, 5, 6, and 7.

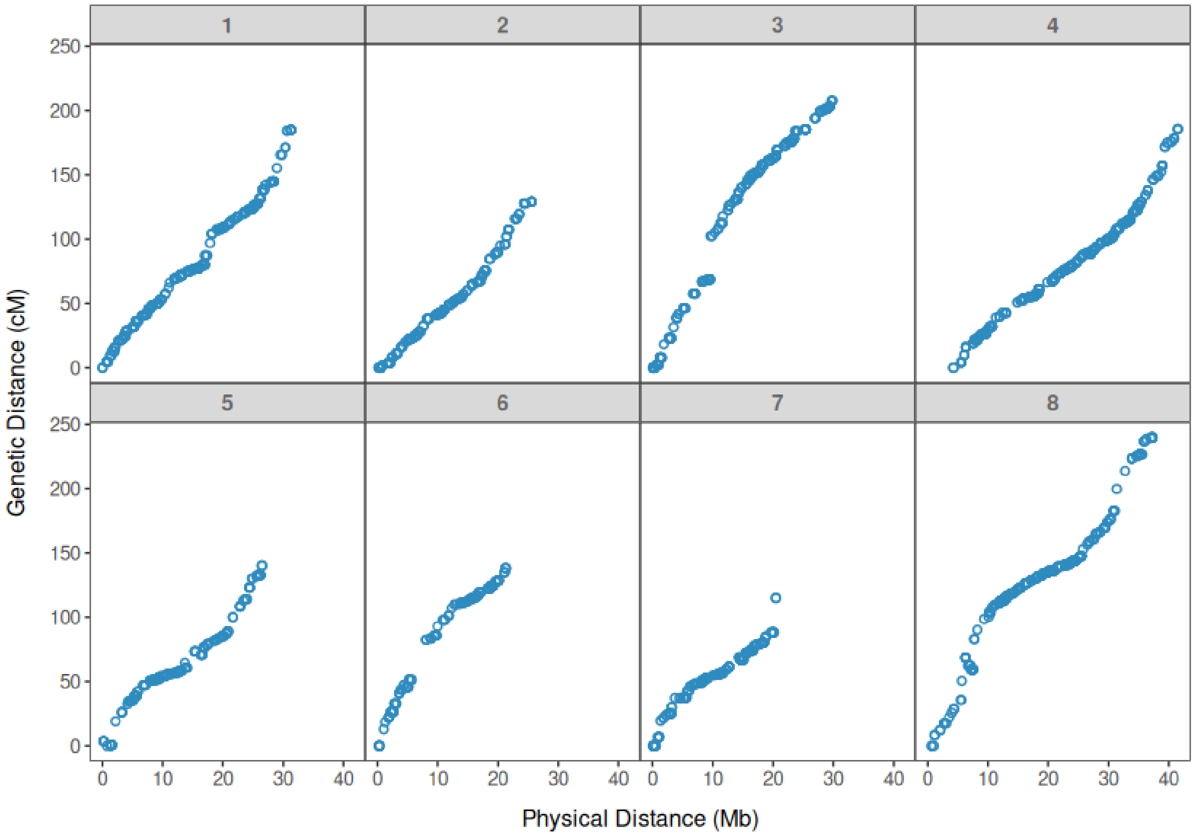


**Figure S6. Correlation between genetic and physical distance of SNPs.** The relationship between genetic and physical distances is shown across the eight linkage groups constructed from 742 F2 individuals genotyped at 2,082 reliable SNP markers. The x-axis represents the physical distance (Mb); the y-axis shows the genetic distance (cM) of markers along the chromosomes.

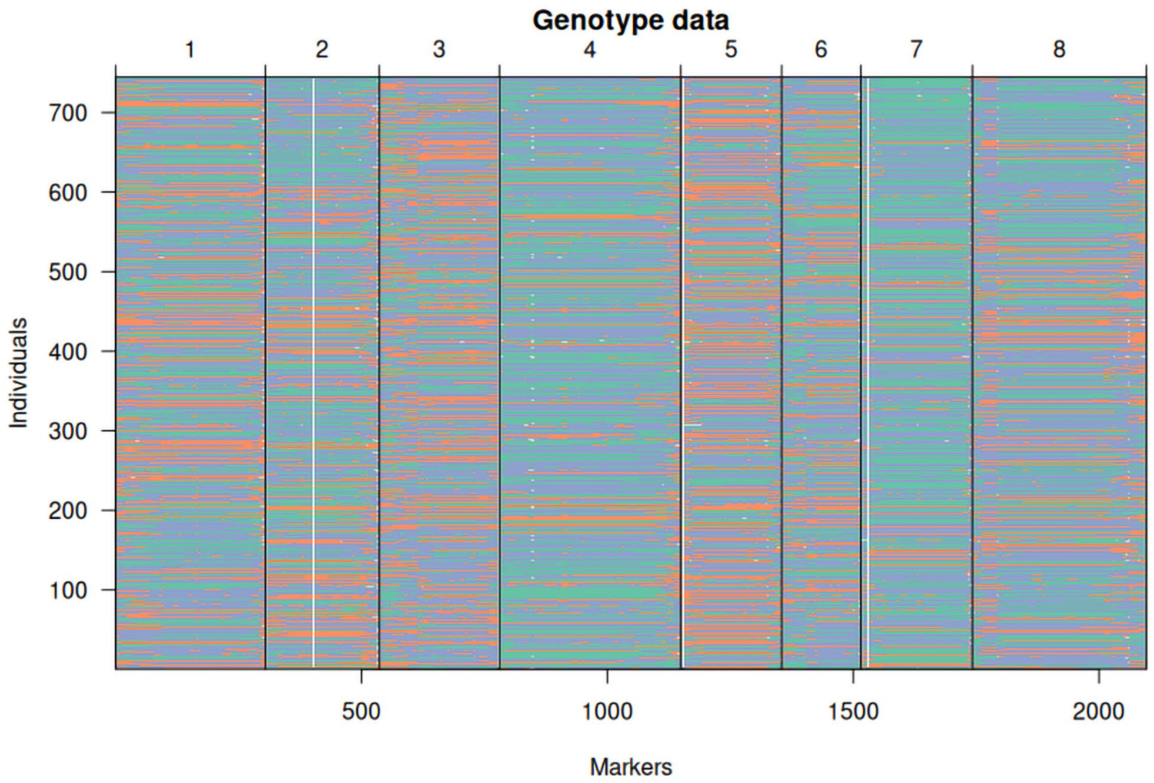


**Figure S7. Mosaic plot of SNP distribution along the genome in the *Arabis* mapping population.** The plot depicts the distribution of 2,082 SNP markers across the genome for each of the 742 individuals in the mapping population. Different colors represent the genotypes observed at each marker: orange = NN, purple = NS, green = SS, and gray = missing data (N: *A. nemorensis* allele; S: *A. sagittata* allele). The y-axis represents individuals, the x-axis shows the SNP markers distributed across 8 chromosomes. This visualization highlights the recombination breakpoints within the chromosomes of each individual.

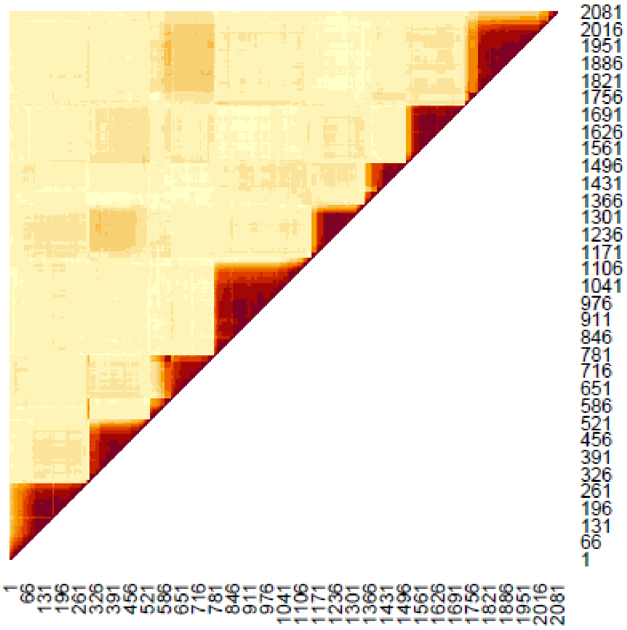


**Figure S8. F2 population linkage disequilibrium.** Linkage disequilibrium in the F2 population quantified as the correlation coefficient of *A. sagittata* allele counts between each pair of markers. The highest linkage is found among loci on the same chromosome, as a result of which the 8 chromosomes are visible and displayed in order. Significant interchromosomal associations of parental alleles were detected between chromosomes 2 and 5, and chromosomes 3 and 8. The resolution of the map is not sufficient to isolate interacting loci. An anomaly is visible on chromosome 3 around the locus showing a strong depletion in heterozygous individuals.

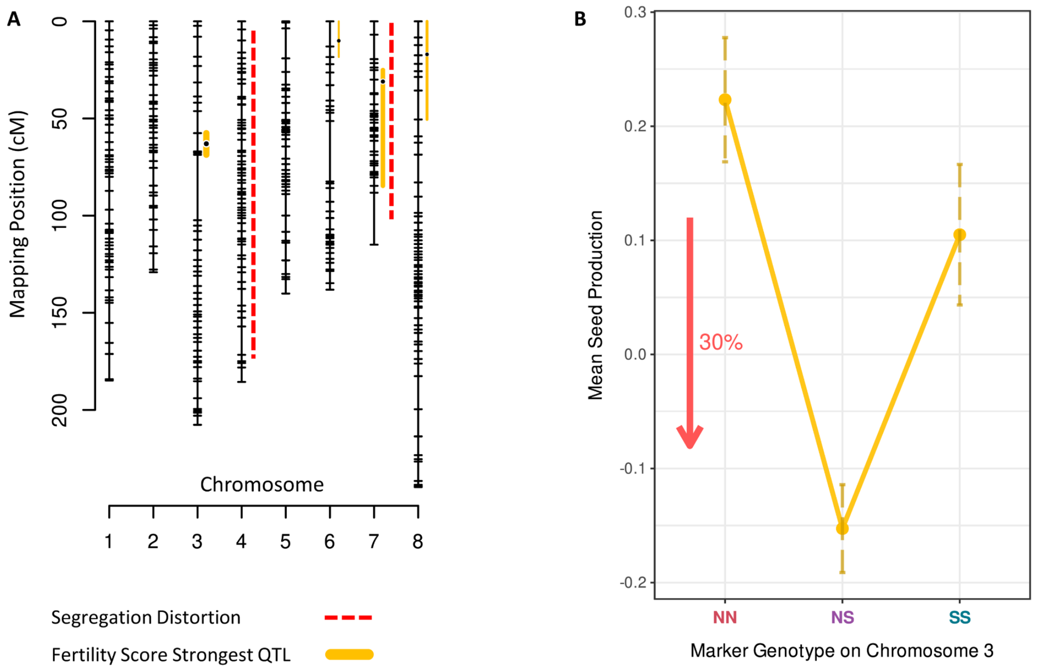


**Figure S9. Genetic architecture of fertility score.** (A) The genetic architecture of Fertility Score and segregation distortion are displayed. The width of the bars represents the strength of the LOD score for each QTL. (B) The effect of the strongest Fertility Score QTL located on chromosome 3 is shown, highlighting its inter-allelic incompatibility.

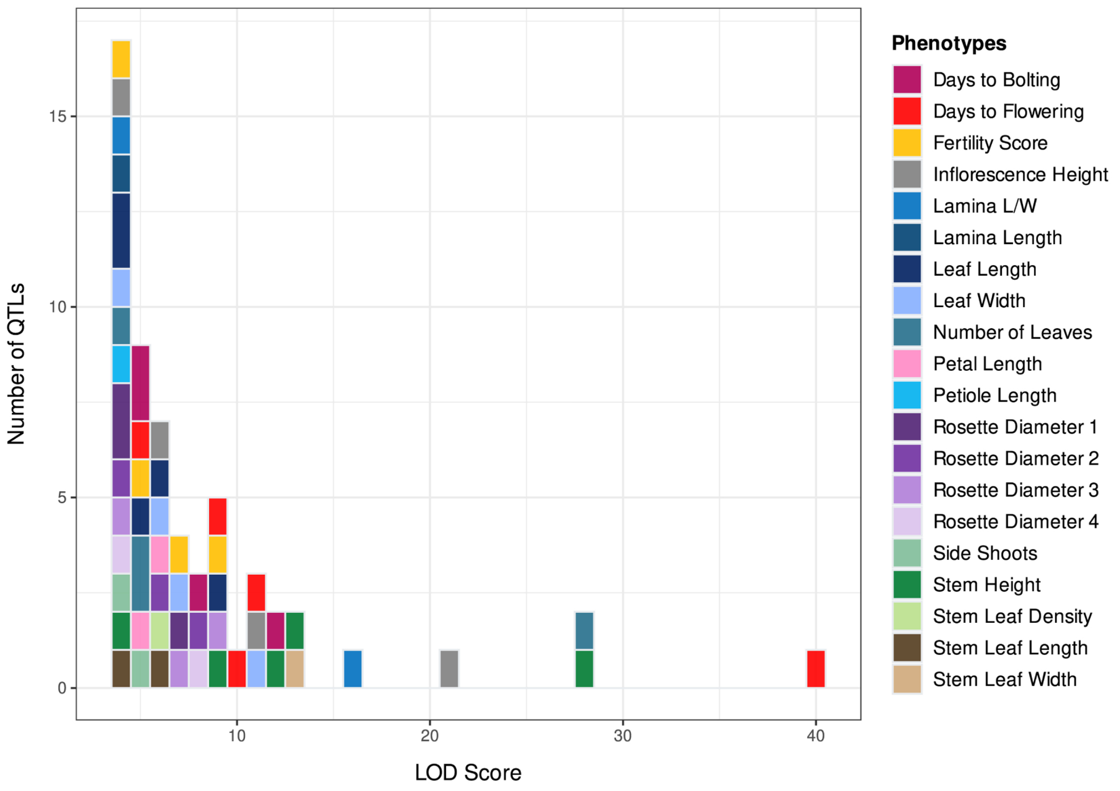


**Figure S10. QTLs and LOD score distribution.** The figure above shows the distribution of ecologically relevant traits in QTLs detected in the F2 population LOD scores. Each block represents one QTL and color phenotypes.

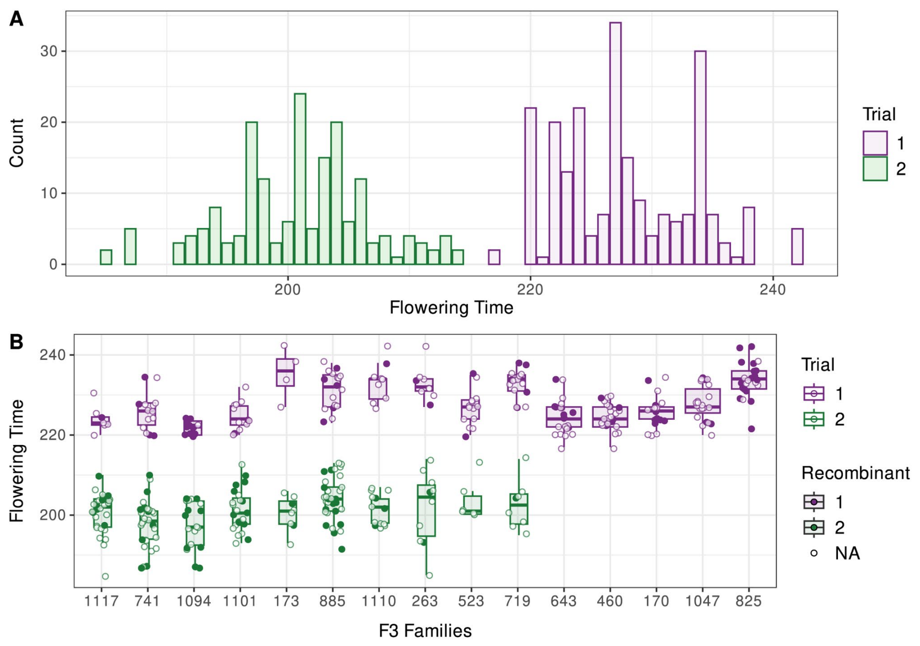


**Figure S11. Distribution of flowering time in *Arabis* F3 hybrids.** The figure represents the distribution of flowering time among 410 F3 plants across two trials. These plants belong to 15 different genotypic F2 families. Individuals with recombination (full dot) allow the QTL region to be narrowed down. In trial 1, family 825, which had the most replicates, had the latest mean flowering time, 233 days (n=23, SD=4.50). Conversely, family 1,094 flowered earliest and had the lowest variation, averaging 222 days (n=15, SD=1.62). In trial 2, plants generally displayed earlier flowering than plants in trial 1 (Table 5). This may be due to differences in environmental conditions such as temperature or light intensity, despite efforts to maintain consistent settings in both common garden experiments. Family 1,094 had the earliest flowering time, with a mean of 198 days (n=19, SD=5.86), whereas family 885 had the latest flowering time,204 days (n=34, SD=5.43). The lowest variation was observed in family 173, the highest in family 263.

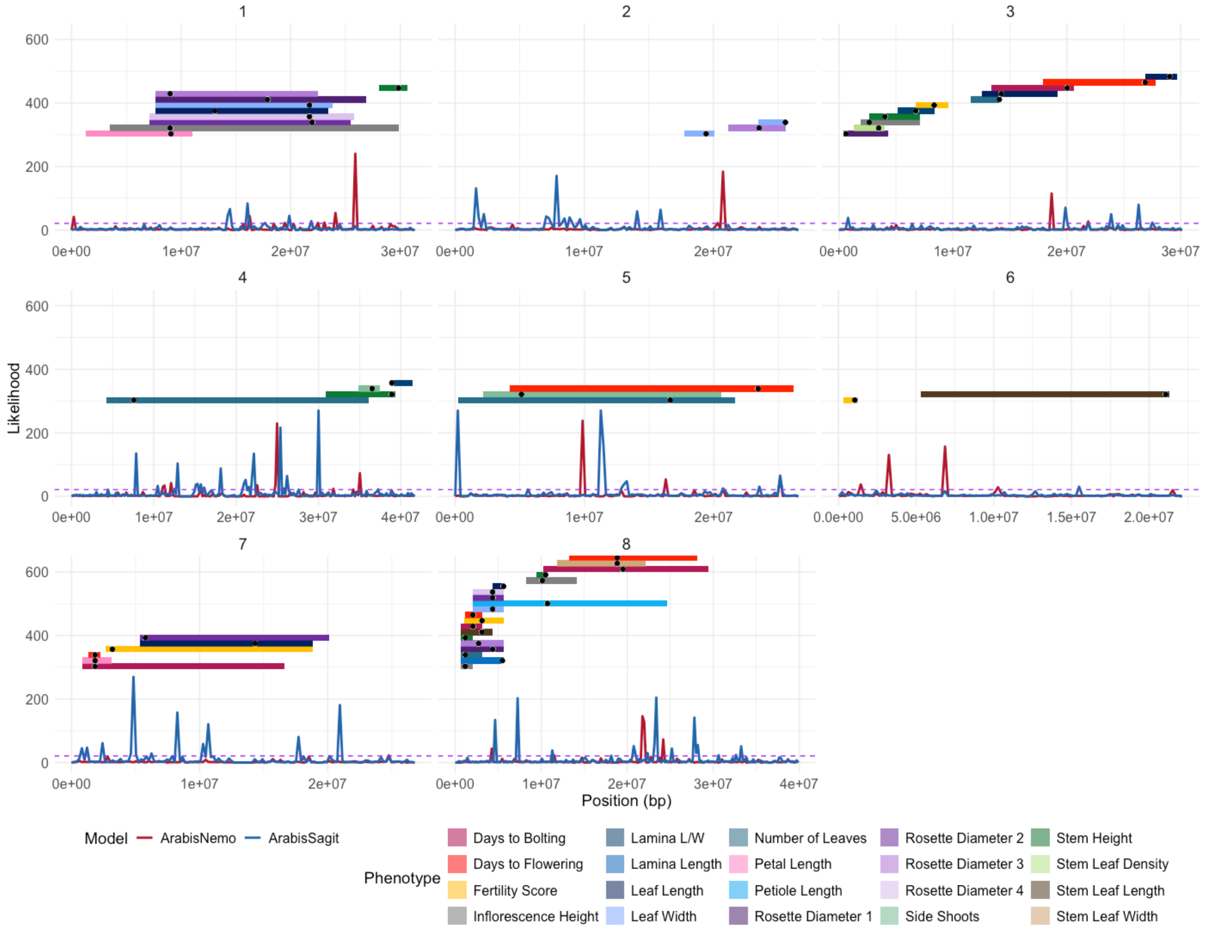


**Figure S12. Sweep detection and QTLs across chromosomes for *A. nemorensis* and *A. sagittata*.** The plot shows detected selective sweeps (lines) and the positions of all QTL regions (rectangles) for various phenotypic traits, with QTLs categorized by trait type and represented in different colors. Each chromosome is displayed in a separate facet, with likelihood values of sweeps plotted along the y-axis. A dashed horizontal line indicates the likelihood threshold for significant sweep detection. Each QTL bar is adjusted in width and positioned along the y-axis based on its chromosome index, and black dots mark QTL peak positions. The likelihood curves are represented for both species, *A. nemorensis* (red) and *A. sagittata* (blue).

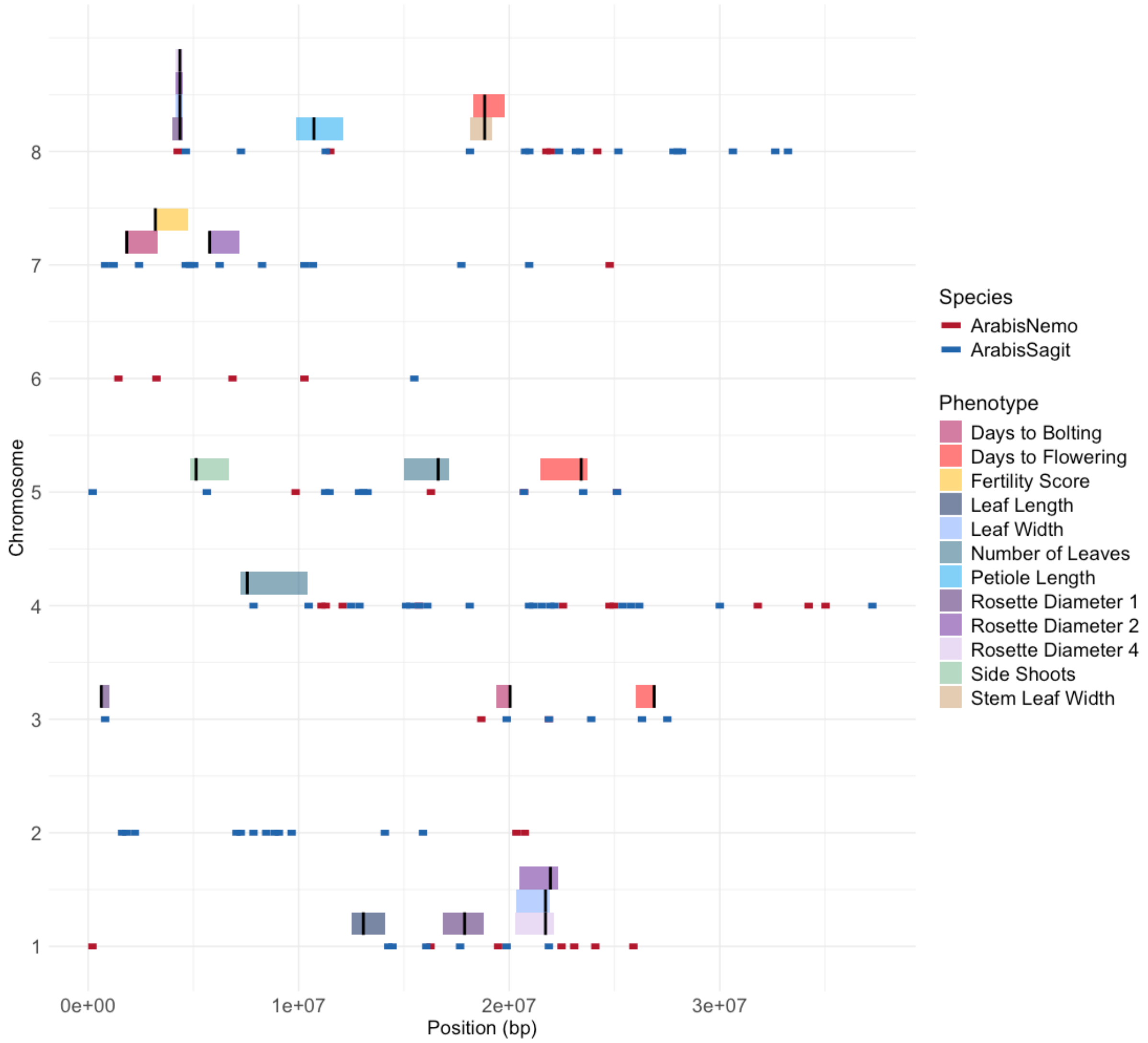


**Figure S13. Overlap between selective sweep windows and 10% quantile QTL regions across chromosomes in *A. nemorensis* and *A. sagittata*.** The plot shows selective sweeps as horizontal lines, each representing a ±100,000 bp region centered around detected sweep positions. Only overlapping QTLs (with sweeps) are shown as colored horizontal bars, extending to a 10% quantile range around their peak positions. Black vertical lines indicate QTL peak positions within these regions. Different colors denote specific phenotypic traits associated with each QTL; sweeps are colored according to regions found within each species.

**Table S1. Results of reciprocal cross-effect analysis on phenotypic traits**. The table summarizes the *p*-values from the analysis of cross-direction and maternal influence on phenotypic traits in the F2 population. Significant *p*-values are bold for emphasis.

| **Trait** | **Estimated Effect *A. sagitatta* Female** | ***p-value*** |
|---|---|---|
| Days to Bolting (B.T) | 0.007349 | 0.59110 |
| Days to Flowering (F.T) | 0.008801 | 0.45541 |
| Fertility Score (W.S) | 0.046190 | 0.85410 |
| Inflorescence Height (P.H) | 0.007968 | 0.91211 |
| Lamina Length (Lam) | -0.120190 | 0.16244 |
| Lamina L/W (LLW) | 0.015500 | 0.75859 |
| Leaf Length (L.L) | -0.043710 | 0.60346 |
| Leaf Width (L.W) | -0.128730 | 0.09654 |
| Number of Stem Leaves (N.L) | -0.054995 | 0.46388 |
| Petal Length (Pet) | 0.015750 | 0.67500 |
| **Petiole Length (Pti)** | **0.681090** | **0.00591** |
| **Rosette Diameter 1 (RD1)** | **-0.323510** | **0.00008** |
| **Rosette Diameter 2 (RD2)** | **-0.207410** | **0.01358** |
| **Rosette Diameter 3 (RD3)** | **-0.193682** | **0.03091** |
| Rosette Diameter 4 (RD4) | -0.000311 | 0.99680 |
| Side Shoots (Ssh) | -0.322387 | 0.05804 |
| Stem Height (S.H) | -0.005128 | 0.93638 |
| Stem Leaf Density (SLD) | -0.023045 | 0.73420 |
| **Stem Leaf Length (SLL)** | **0.215120** | **0.00003** |
| Stem Leaf Width (SLW) | 0.066639 | 0.27150 |
| **Petiole L/Lamina L (P.L)** | **0.698930** | **0.01190** |
| Ground Shoots (Gsh) | -0.223140 | 0.47830 |

**Table S2. Genetic map overview.** This table summarizes the physical and genetic lengths of each chromosome, along with the number of SNP markers. The total number of SNPs included in the map is 2,082.

| Chromosome | Physical Length (Mb) | Genetic Length (cM) | Number of SNPs |
|---|---|---|---|
| 1 | 32 | 185 | 304 |
| 2 | 26 | 130 | 232 |
| 3 | 30 | 208 | 244 |
| 4 | 41 | 186 | 369 |
| 5 | 27 | 140 | 202 |
| 6 | 22 | 138 | 160 |
| 7 | 21 | 115 | 220 |
| 8 | 38 | 240 | 351 |

**Table S3. Overview of mean flowering time for *Arabis* F3 families.** This table provides the mean flowering time for each F3 family across the two trials of the fine-mapping experiment. Trial 2 did not include all 15 families.

| F3 Family | Mean Flowering Time in Trial 1 | Mean Flowering Time in Trial 2 |
|---|---|---|
| 170 | 226 | *NA* |
| 173 | 235 | 200 |
| 263 | 233 | 202 |
| 460 | 225 | *NA* |
| 523 | 227 | 204 |
| 643 | 224 | *NA* |
| 719 | 233 | 202 |
| 741 | 226 | 198 |
| 825 | 234 | *NA* |
| 885 | 231 | 204 |
| 1047 | 228 | *NA* |
| 1094 | 222 | 198 |
| 1101 | 225 | 201 |
| 1110 | 233 | 201 |
| 1117 | 224 | 201 |

**Table S4. Genotype and phenotype of F3 families used in the flowering time fine-mapping experiment.** The table below shows the genotypes for all flowering time QTLs. Q4 is the strongest QTL which was used for fine mapping. “S” represents *A. sagittata* homozygous; “N,” *A. nemorensis* homozygous; and “SN,” heterozygous genotypes. The flowering time observed in the F2 common garden experiment is also listed. Family 173 is not included in this table. These families were selected based on a genetic map constructed using an earlier version of the genome assembly.

| QTL ID | Chr | 170 | 263 | 460 | 523 | 643 | 719 | 741 | 825 | 885 | 1047 | 1094 | 1101 | 1110 | 1117 |
|---|---|---|---|---|---|---|---|---|---|---|---|---|---|---|---|
| **Q1** | 3 | N | S | N | S | N | S | N | S | N | N | N | N | S | N |
| **Q2** | 5 | N | S | S | NS | N | N | N | NS | N | N | S | N | NS | NS |
| **Q3** | 7 | N | NS | S | N | S | S | NS | S | S | S | NS | NS | NS | NS |
| **Q4** | 8 | NS | NS | NS | NS | NS | NS | NS | NS | NS | NS | NS | NS | NS | NS |
| **Q5** | 8 | N | S | N | S | N | S | N | S | S | NS | N | N | S | S |
| **Flowering time** | | 190 | 196 | 187 | 193 | NA | 195 | NA | 195 | 193 | 183 | 181 | 181 | NA | 183 |

**Table S5. Summary of detected QTLs across traits and their relationship to fertility and distortion regions.** This table includes the trait name, chromosome (Chr), QTL peak position (in bp and cM), and the corresponding LOD score for each detected QTL. The final column ("Independence") indicates whether each QTL is classified as independent (TRUE) or not independent (FALSE), based on its location and potential overlap with fertility-associated QTLs or known segregation distortion regions (chromosomes 4 and 7). QTLs were considered independent if they were not located on chromosomes 4 or 7 and did not overlap with any fertility QTLs. In total, 48.3% of the QTLs were classified as independent.

| Trait | QTL ID | Chr | Position (bp) | Position (cM) | LOD | Independence |
|---|---|---|---|---|---|---|
| Days to Bolting | Q1 | 3 | 20040893 | 163.00 | 5.15 | TRUE |
| Days to Bolting | Q2 | 7 | 1835566 | 21.49 | 5.36 | FALSE |
| Days to Bolting | Q3 | 8 | 2061023 | 12.24 | 7.79 | FALSE |
| Days to Bolting | Q4 | 8 | 19500950 | 134.21 | 12.40 | TRUE |
| Days to Flowering | Q1 | 3 | 26882840 | 190.00 | 10.00 | TRUE |
| Days to Flowering | Q2 | 5 | 23418119 | 111.00 | 4.69 | TRUE |
| Days to Flowering | Q3 | 7 | 1835566 | 21.49 | 11.23 | FALSE |
| Days to Flowering | Q4 | 8 | 2061023 | 13.00 | 40.45 | FALSE |
| Days to Flowering | Q5 | 8 | 18832553 | 133.00 | 8.91 | TRUE |
| Fertility Score | Q1 | 3 | 8362080 | 63.00 | 8.77 | FALSE |
| Fertility Score | Q2 | 6 | 1052484 | 10.00 | 3.90 | FALSE |
| Fertility Score | Q3 | 7 | 3192272 | 31.00 | 6.66 | FALSE |
| Fertility Score | Q4 | 8 | 3118527 | 17.00 | 5.07 | FALSE |
| Inflorescence Height | Q1 | 1 | 9004320 | 49.89 | 4.17 | TRUE |
| Inflorescence Height | Q2 | 3 | 2664384 | 27.00 | 11.39 | FALSE |
| Inflorescence Height | Q3 | 8 | 1180103 | 6.00 | 20.79 | FALSE |
| Inflorescence Height | Q4 | 8 | 10128705 | 101.00 | 6.38 | TRUE |
| Lamina Length | Q1 | 8 | 5481058 | 36.00 | 4.33 | FALSE |
| Lamina L/W | Q1 | 3 | 6744303 | 61.00 | 15.64 | FALSE |
| Lamina L/W | Q2 | 4 | 38897287 | 163.00 | 4.04 | FALSE |
| Leaf Length | Q1 | 1 | 13071968 | 72.39 | 4.46 | TRUE |
| Leaf Length | Q2 | 3 | 14255522 | 136.22 | 5.21 | TRUE |
| Leaf Length | Q3 | 3 | 29031007 | 201.41 | 5.62 | TRUE |
| Leaf Length | Q4 | 7 | 14342773 | 68.78 | 4.28 | FALSE |
| Leaf Length | Q5 | 8 | 5624776 | 35.00 | 9.43 | FALSE |
| Leaf Width | Q1 | 1 | 21722429 | 116.00 | 3.87 | TRUE |

| Trait | QTL ID | Chr | Position (bp) | Position (cM) | LOD | Independence |
|---|---|---|---|---|---|---|
| Leaf Width | Q2 | 2 | 19435694 | 87.00 | 7.10 | TRUE |
| Leaf Width | Q3 | 2 | 25576894 | 129.00 | 6.34 | TRUE |
| Leaf Width | Q4 | 8 | 4356973 | 32.00 | 10.78 | FALSE |
| Number of Leaves | Q1 | 3 | 14075550 | 130.95 | 4.58 | TRUE |
| Number of Leaves | Q2 | 4 | 7558305 | 18.00 | 4.00 | FALSE |
| Number of Leaves | Q3 | 5 | 16624270 | 70.69 | 5.38 | TRUE |
| Number of Leaves | Q4 | 8 | 1180103 | 7.00 | 28.11 | FALSE |
| Petal Length | Q1 | 1 | 9074627 | 51.00 | 4.86 | TRUE |
| Petal Length | Q2 | 7 | 1835566 | 21.49 | 5.69 | FALSE |
| Petiole Length | Q1 | 8 | 10723493 | 107.50 | 3.81 | FALSE |
| Rosette Diameter 1 | Q1 | 1 | 17879753 | 95.00 | 4.50 | TRUE |
| Rosette Diameter 1 | Q2 | 3 | 623617 | 2.31 | 3.63 | TRUE |
| Rosette Diameter 1 | Q3 | 8 | 4362085 | 28.00 | 6.94 | FALSE |
| Rosette Diameter 2 | Q1 | 1 | 21954996 | 115.00 | 6.16 | TRUE |
| Rosette Diameter 2 | Q2 | 7 | 5769578 | 42.70 | 3.84 | FALSE |
| Rosette Diameter 2 | Q3 | 8 | 4356973 | 31.00 | 7.82 | FALSE |
| Rosette Diameter 3 | Q1 | 1 | 9004320 | 49.89 | 9.39 | TRUE |
| Rosette Diameter 3 | Q2 | 2 | 23541930 | 118.00 | 4.02 | TRUE |
| Rosette Diameter 3 | Q3 | 8 | 2718077 | 18.00 | 7.21 | FALSE |
| Rosette Diameter 4 | Q1 | 1 | 21722429 | 115.29 | 4.33 | TRUE |
| Rosette Diameter 4 | Q2 | 8 | 4356973 | 32.00 | 7.81 | FALSE |
| Side Shoots | Q1 | 4 | 36501835 | 137.73 | 3.86 | FALSE |
| Side Shoots | Q2 | 5 | 5129643 | 35.50 | 4.88 | TRUE |
| Stem Height | Q1 | 1 | 29847516 | 162.00 | 4.37 | TRUE |
| Stem Height | Q2 | 3 | 4035375 | 36.00 | 13.09 | FALSE |
| Stem Height | Q3 | 4 | 38897287 | 160.00 | 8.78 | FALSE |
| Stem Height | Q4 | 8 | 1180103 | 6.00 | 28.31 | FALSE |

| Trait | QTL ID | Chr | Position (bp) | Position (cM) | LOD | Independence |
|---|---|---|---|---|---|---|
| Stem Height | Q5 | 8 | 10506892 | 102.00 | 12.25 | TRUE |
| Stem Leaf Density | Q1 | 3 | 3491752 | 29.00 | 5.76 | TRUE |
| Stem Leaf Length | Q1 | 6 | 21070306 | 134.77 | 5.92 | TRUE |
| Stem Leaf Length | Q2 | 8 | 3118527 | 15.00 | 3.97 | FALSE |
| Stem Leaf Width | Q1 | 8 | 18832576 | 132.55 | 13.23 | TRUE |
